# Spontaneous body wall contractions stabilize the fluid microenvironment that shapes host-microbe associations

**DOI:** 10.1101/2022.11.30.518486

**Authors:** JC Nawroth, C Giez, A Klimovich, E Kanso, TCG Bosch

## Abstract

The freshwater polyp Hydra is a popular biological model system; however, we still do not understand one of its most salient behaviours, the generation of spontaneous body wall contractions. Here, by applying experimental fluid dynamics analysis and mathematical modelling, we provide functional evidence that spontaneous contractions of body walls enhance the transport of chemical compounds from and to the tissue surface where symbiotic bacteria reside. Experimentally, a reduction in the frequency of spontaneous body wall contractions is associated with a changed composition of the colonizing microbiota. Together, our findings suggest that spontaneous body wall contractions create an important fluid transport mechanism that (1) may shape and stabilize specific host-microbe associations and (2) create fluid microhabitats that may modulate the spatial distribution of the colonizing microbes. This mechanism may be more broadly applicable to animal-microbe interactions since research has shown that rhythmic spontaneous contractions in the gastrointestinal tracts are essential for maintaining normal microbiota.

## INTRODUCTION

The millimeter-scale freshwater polyp Hydra with its simple tube-like body structure (Fig. 1A) and stereotypic movement patterns is a popular biological model organism for immunology [1; 2; 3; 4; 5], developmental and evolutionary biology, and neurobiology [4; 6]. Despite Hydra’s relevance for fundamental research, one of the animal’s most salient behaviours remains a mystery since its first description in 1744 [7]: it is unclear why Hydra undergoes recurrent full body contraction-relaxation cycles, approximately one every 10 - 30 minutes (Fig. 1B), while attached to the substratum. Most, if not all, animals exhibit such spontaneous contractions (SCs) of muscular organs and body walls to pump internal fluids, feed, locomote, or clean surfaces [8; 9; 10; 11; 12]. However, Hydra’s morphology and SC characteristics do not fit any of these common functions. Here, we sought to investigate the functional underpinnings of Hydra’s SCs from a fluid mechanics perspective. We hypothesized that the SCs might generate fluid transport patterns of relevance to Hydra’s microbial partners that colonize the so-called glycocalyx on the outer body wall [5; 13]. (Fig. 1C). Typically, the composition and distribution of symbiotic microbial communities residing at solid-liquid interfaces, such as in the gut and lung, are regulated both by interaction with host cells and by the properties of the fluid medium [14; 15]. Therefore, we asked whether Hydra’s microbiome might be influenced by fluid flow associated with SCs.

**Figure 1:**
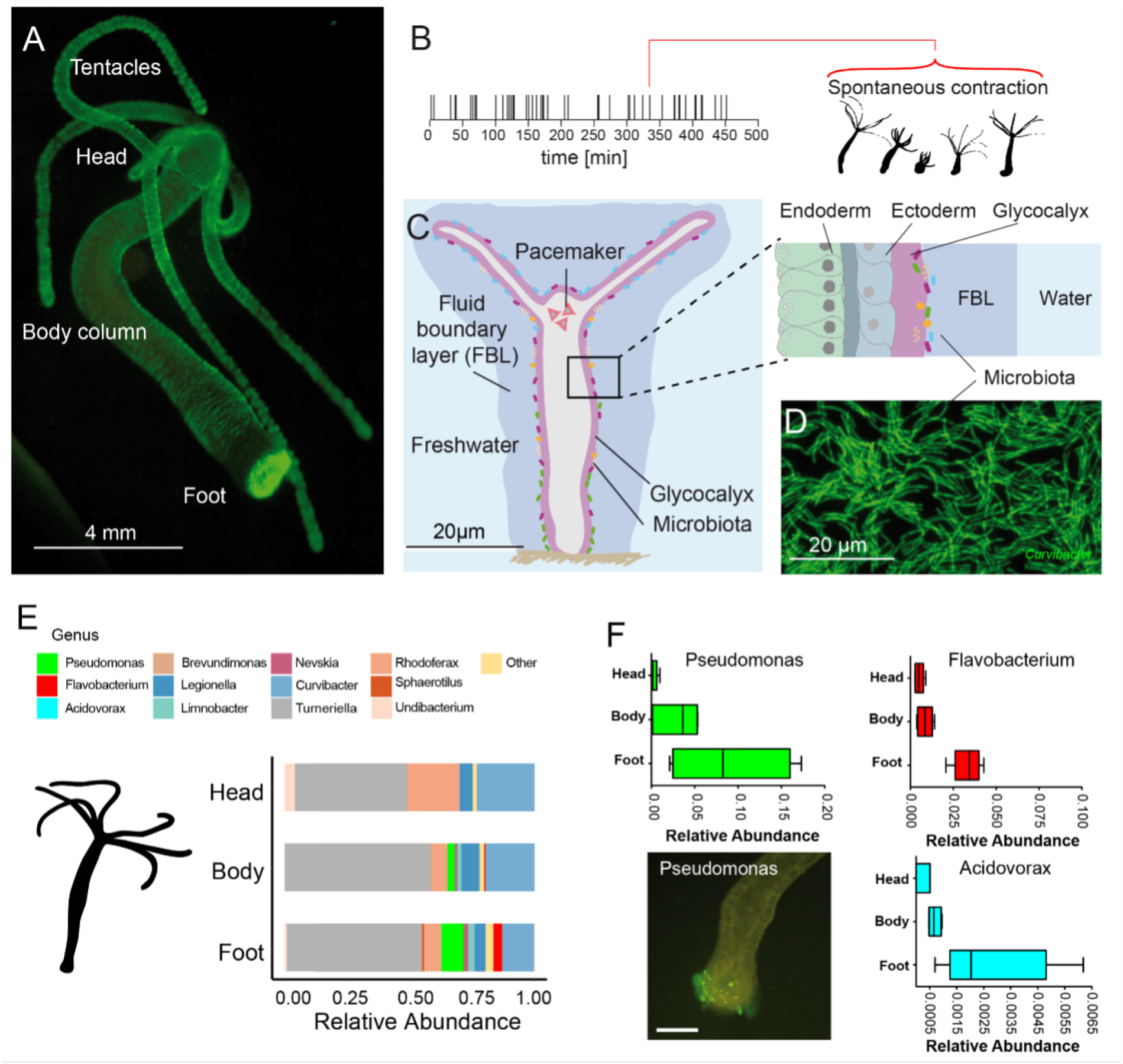
The freshwater polyp Hydra, a model system for the role of spontaneous body wall contractions in shaping microbe biogeography. **A**. Polyp colonized with fluorescent labelled betaproteobacteria. **B**. A representative contractile activity pattern of an individual polyp recorded over 8 h (left). Each dash on the timeline represents an individual spontaneous contraction-relaxation cycle (right). **C**. Schematic representation of a Hydra with a fluid boundary layer (FBL) surrounding the polyp with the glycocalyx layer adjacent to the polyp’s tissue. Inset: Tissue architecture covered outside by the mucus-like glycocalyx which provides the habitat for a specific bacterial community on the interface with the FBL. **D**. Dense community of fluorescently labelled bacteria (main colonizer Curvibacter spec.) colonizing the polyp’s head. **E**. The biogeography of Hydras symbionts under undisturbed/control conditions follows a distinct spatial colonization pattern along the body column. **F**. Bacteria mostly colonizing the foot region include Pseudomonas, Flavobacterium, and Acidovorax, scalebar: 500 *μ*m.

## RESULTS

First, we mapped the abundance and distribution of microbiota that reside on the glycocalyx (Fig. 1C, D). The glycocalyx is a multi-layered extracellular cuticle covering Hydra’s ectodermal epithelial cells, has mucus-like properties, and shapes the microbiome by providing food and antimicrobial peptides [15]. Regional differences in the glycocalyx have not been reported; however, using 16S rRNA profiling on different sections of Hydra, we found that different microbial species favor different regions along the body column, including foot- and head-dominating microbiota (Fig. 1E and F). Based on these intriguing data and recent insights into how fluid flow shapes spatial distributions of bacteria [16], we hypothesized that the SCs might generate fluidic microhabitats that facilitate the microbial biogeography along the body column.

### Kinematics and flow physics of individual spontaneous contractions

To explore this hypothesis, we used video microscopy and analyzed the animal’s body kinematics to assess the differential impact of spontaneous contractions on the fluid microenvironment. We examined the contraction activity (Fig. 2A) of multiple animals over 8 hours and we recorded an average contraction frequency of 2.5 spontaneous contractions per hour (CPH) (Suppl. Fig. 1A, control condition), corresponding to an average duration (*T*_IC_) of the inter-contraction intervals (IC) of 24 min. To arrive at estimates rooted in probability theory that take into consideration the distribution of *T*_IC_, we collected the measured *T*_IC_ from all animals in the form of a histogram (Suppl. Fig. 2A, control condition). The resulting distribution of *T*_IC_ is best fit by an exponential distribution, implying that contraction events of individual polyps follow a stochastic Poisson process. We computed the average frequency based on the exponential fit to these biological data. We found that the so-computed contraction frequency is equal to 2.9 CPH, which is slightly larger than the 2.5 CPH obtained by taking a direct average of the data.

**Figure 2:**
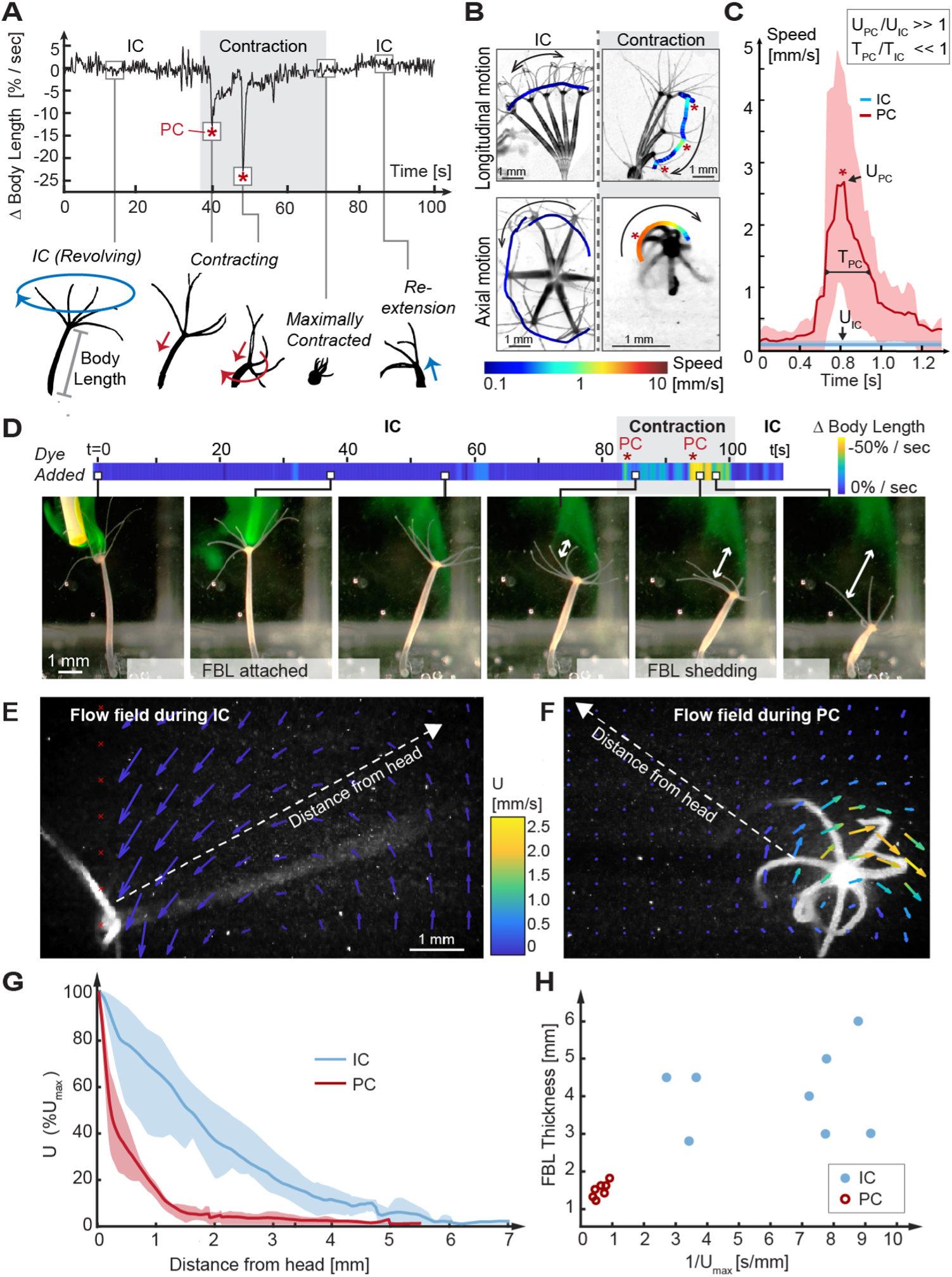
Kinematic and fluid dynamics analysis of individual contraction events reveal the shedding of the fluid boundary layer during each spontaneous contraction event. **A**. Top: Representative plot of the change in relative body length in Hydra as a function of time shows transition from an IC to a SC with two peak contraction events (PC, asterisks), and the return to IC. Bottom: Typical kinematic pattern associated with ICs and contractions. Arrows indicate distinct body trajectories during ICs (blue) compared to contractions (red). **B**. Typical body trajectories in longitudinal and axial plane during ICs and SCs visualized by time-lapse microscopy. Maximal speeds (log10 scale) are indicated by color-coded trajectories of the head (oral end or tentacles). Trajectories are slightly offset to avoid obscuring the animal. Asterisks denote PC events. **C**. Comparison of maximal velocities near head reached during IC (n=3 animals) and PC (n=5 animals). Lines: average curves. Shaded areas: interquartile range. Inset: Relative scaling of PC duration (*T*_PC_), IC duration (*T*_IC_), PC velocity magnitudes (*U*_PC_) and IC velocity magnitudes (*U*_IC_). **D**. Application of a fluorescent dye reveals existence of a fluid boundary layer during IC and its shedding upon a contraction event. A representative time-lapse series. White arrows indicate FBL shedding after PCs, i.e., the growing separation between the original, stained FBL and Hydra’s head. **E**. Quantification of Hydra’s flow velocity field during IC and F. during a typical PC with axial rotation (top view). Flow vectors and velocities are indicated by color-coded arrows. **G**. Relative change of fluid flow speed as a function of distance from Hydra’s surface (measured along dotted lines in **E, F**) during IC (blue) and during PCs with rotation (red). Lines: average curves. Shaded areas: interquartile range. **H**. FBL thickness, defined as distance from Hydra at which 90% freestream speed is reached, is inversely correlated to maximal flow speed (*U*_max_).

Each stage of the IC and SC cycle is characterized by stereotypic kinematic patterns of Hydra’s motion trajectories and velocities (Fig. 2A, bottom). During IC periods, which on average last tens of minutes (Fig. 2A, Suppl. Fig. 2A control condition), Hydra’s body column remains nearly fully extended, and slowly revolves at full length around its foothold, tracing a cone-shaped volume over time (Fig. 2B, Suppl. Video 1). By contrast, SC events typically last a few seconds and are characterized by a stepwise shortening of the body column with one or more peak contraction events until the body column has shortened to 20% or less of its original length (Fig. 2A-C). A peak contraction (PC) is defined as any part of the SC during which Hydra contracts at a rate of 25% change in body length per second or more. SCs are frequently accompanied by axial rotation in a spiralling downward motion (Fig. 2A and B, Suppl. Video 1) and are typically followed by slow re-extension to IC length (Fig. 2A) in a random direction (Suppl. Fig. 3). We quantified the speeds and timescales that different regions of Hydra’s body column experience during ICs and SCs. During SCs, specifically during PCs characterized by simultaneous linear shortening and axial rotation, the oral region accelerates to peak velocities *U*_PC_ of 10 mm/s. This is almost two orders of magnitude faster than maximal head speeds *U*_IC_ during ICs, which are on the order of 0.1 mm/s in both longitudinal and axial directions (Fig. 2B and C). Further, PCs rarely last longer than *T*_PC_ = 1 s and hence operate at 1000-fold smaller timescales than ICs (with mean duration of ca. 24 minutes, i.e., 1440 seconds).

**Figure 3:**
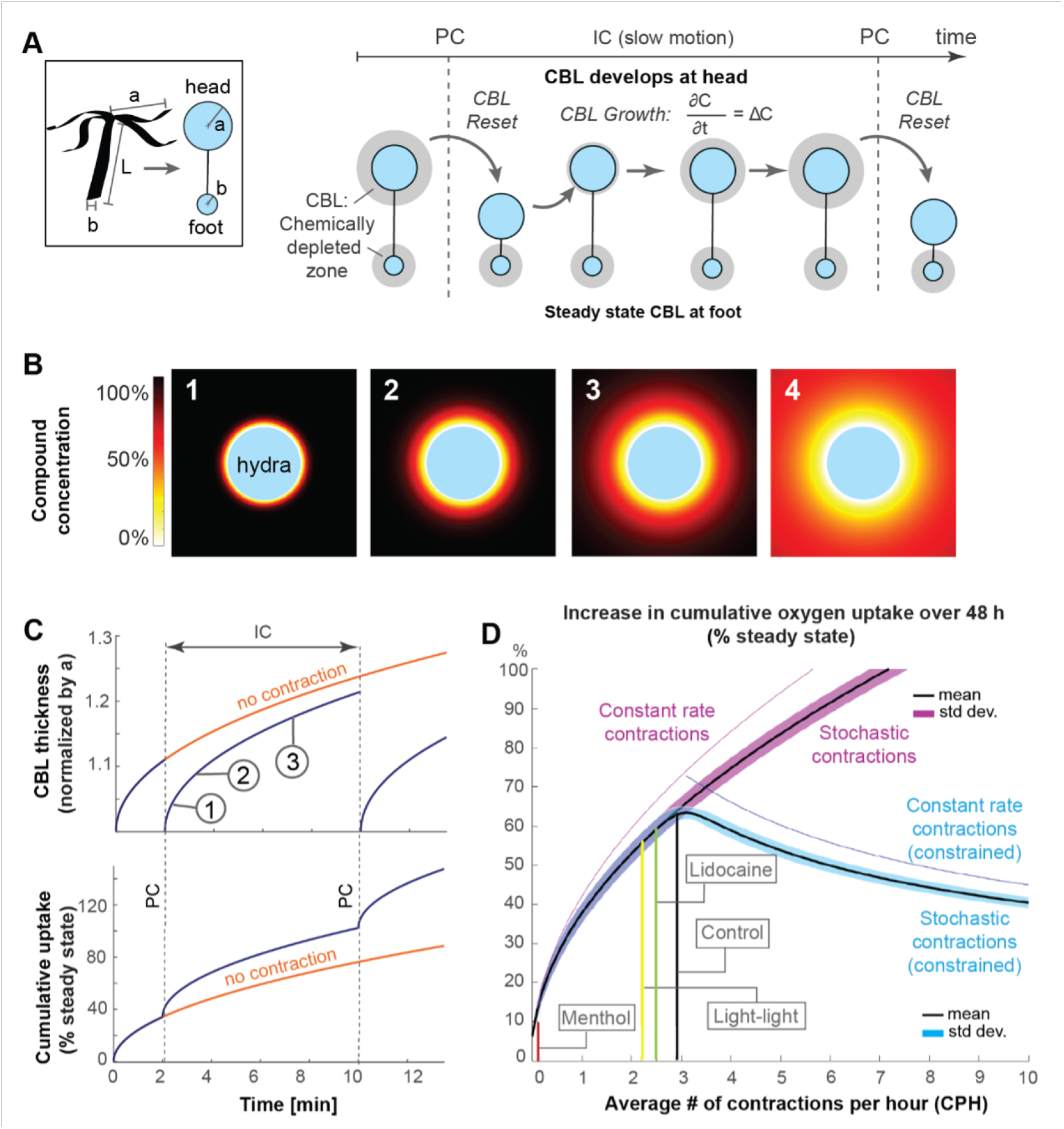
Mathematical model suggests that spontaneous contractions enhance mass transport to and from the surface. **A**. Left, box: Simplified animal geometry assumed in model consists of a sphere representing the head and a smaller sphere representing the basal foot. Right: By Fick’s law, a chemically depleted concentration boundary layer (CBL) forms around Hydra’s head and foot region through continuous uptake of chemical compounds at the surface. The CBL is shed from the head region, but not the foot region, through contractions. **B**. Computationally modelled growth of chemically depleted CBL around Hydra’s head over time until steady state is reached (panel 4), assuming continuous uptake at surface and unlimited supply at far distance. **C**. Top: Growth of chemically depleted CBL as a function of time with and without contractions. Bottom: Instantaneous and cumulative uptake rate, respectively, of a given chemical compound (here: oxygen) during the CBL dynamics above; **D**. Predicted increase in cumulative uptake rate (as percentage of steady state) of a given chemical compound over 48 h as a function of SC frequencies within the observed range (purple), and of increasing frequencies with unconstraint number of SCs (magenta) compared to a constraint number of SCs (blue graph).

To assess the effects of Hydra’s body kinematics on its fluid environment, we computed the Reynolds number Re= *ℓU/μ* at the tentacles near Hydra’s mouth, where the tentacle diameter is *ℓ* = 0.1 mm, the kinematic viscosity of water at 20°C is *μ* = 1.0034 mm^2^/s, and the tentacle speed *U* is in mm/s. The Re number compares the effects of fluid inertia to viscous drag forces. During IC, *U* = *U*_*IC*_ is on the order of 0.1 mm/s, and Re is on the order of 0.01. At such low Re number, fluid flow is dominated by viscous effects and distinguished by the presence of a laminar and expansive fluid boundary layer (FBL). The fluid boundary layer can be envisioned as an envelope of fluid enclosing Hydra and moving at the same velocity as Hydra’s surface. The thickness of the fluid boundary layer is defined as the distance from the surface at which the fluid speed decreases by 90% compared to Hydra’s speed, i.e., the distance at which the fluid is no longer following Hydra’s surface [17]. At PC, Hydra’s speed *U* = *U*_PC_ is on the order of 10 mm/s, Re increases to the order of 1, which indicates greater inertial effects that result in thinning of the fluid boundary layer [18].

To label the fluid boundary layer and assess the animal-fluid interactions experimentally [19], we added fluorescent dye adjacent to Hydra’s oral region while in IC state. In one representative recording (Fig. 2D, left; Suppl. Video 2), the dye faithfully followed the fully extended animal’s slow revolutions during the IC for more than one minute, qualitatively confirming the presence of a stably attached fluid boundary layer. Intriguingly, as the animal underwent a SC (with two PC events), the dye separated from the animal’s surface and stayed behind as Hydra retracted, indicating shedding of the original fluid boundary layer (Fig. 2D, right; Suppl. Video 2). Since the animals tend to slowly re-extend into a random direction (Suppl. Fig. 3), their chances of re-encountering the shed fluid are small. To confirm these observations quantitatively and to measure the dynamically changing fluid boundary layer thickness, we recorded the microscale fluid motion using particle imaging velocimetry (PIV) as described previously [20; 21]. During ICs, fluid parcels at distances far from the body column followed the animal’s trajectory, reflecting an expansive fluid boundary layer of almost one full body length in thickness (FBL thickness = 5 mm) (Fig. 2E and G; Suppl. Fig. 4A, Suppl. Video 3). In contrast, during PCs, only the fluid close to Hydra’s surface accelerated with the body whereas fluid at further distances remained unaffected (FBL thickness =1.5 mm) (Fig. 2F and G; Suppl. Fig. 4A, Suppl. Video 3). The thickness of the fluid boundary layer correlated inversely with body speed (Fig. 2H). These measurements demonstrate shedding of the fluid boundary layer during contractions and significant reduction of the thickness of the fluid boundary layer that had developed during IC. Thus, recurrent SCs result in regular shedding of the fluid boundary layer (as illustrated by the streak of dye left behind in Fig. 2D), enabling Hydra to partially escape from its previous fluid environment and thereby transiently reshape the chemical microenvironment at the epithelial surface where the microbial symbionts are localized. Importantly, the contraction events form a brief perturbation of the IC flow regime at the oral region, while the foot region remains motionless even during PCs (Fig. 2B) and experiences slow flow speeds of maximal values on the order of 0.1 mm/s (Suppl. Fig. 4B). These flow speeds are comparable to flow speeds experienced at the head between contractions (Suppl. Fig. 4A). Taken together, these results suggest that Hydra’s SCs lead to a maximal shedding of viscous boundary layers near the head’s surface, and minimal shedding near the foot’s surface. This differentiation in the fluid environment could be relevant because of our finding that Hydra’s surface is colonized by foot- and head-dominating microbiota (Fig. 1D and E). Contraction-induced fluid boundary layer shedding may enhance the transport of bacteria-relevant compounds, such as metabolites, antimicrobials, and extracellular vesicles [22; 23], between Hydra’s head and the fluid environment, compared to lower transport near the foot, hence generating biochemical microhabitats which could promote the observed microbial biogeography. This hypothesis is not readily amenable to experimental interrogation. To date, there are no universal tools for experimentally identifying individual and combinations of the many chemical compounds released or absorbed by microbes [24]. We therefore probed this hypothesis indirectly. We developed a simple mathematical model of chemical transport in a fluid environment that gets regularly reset by shedding events, as discussed next.

**Figure 4:**
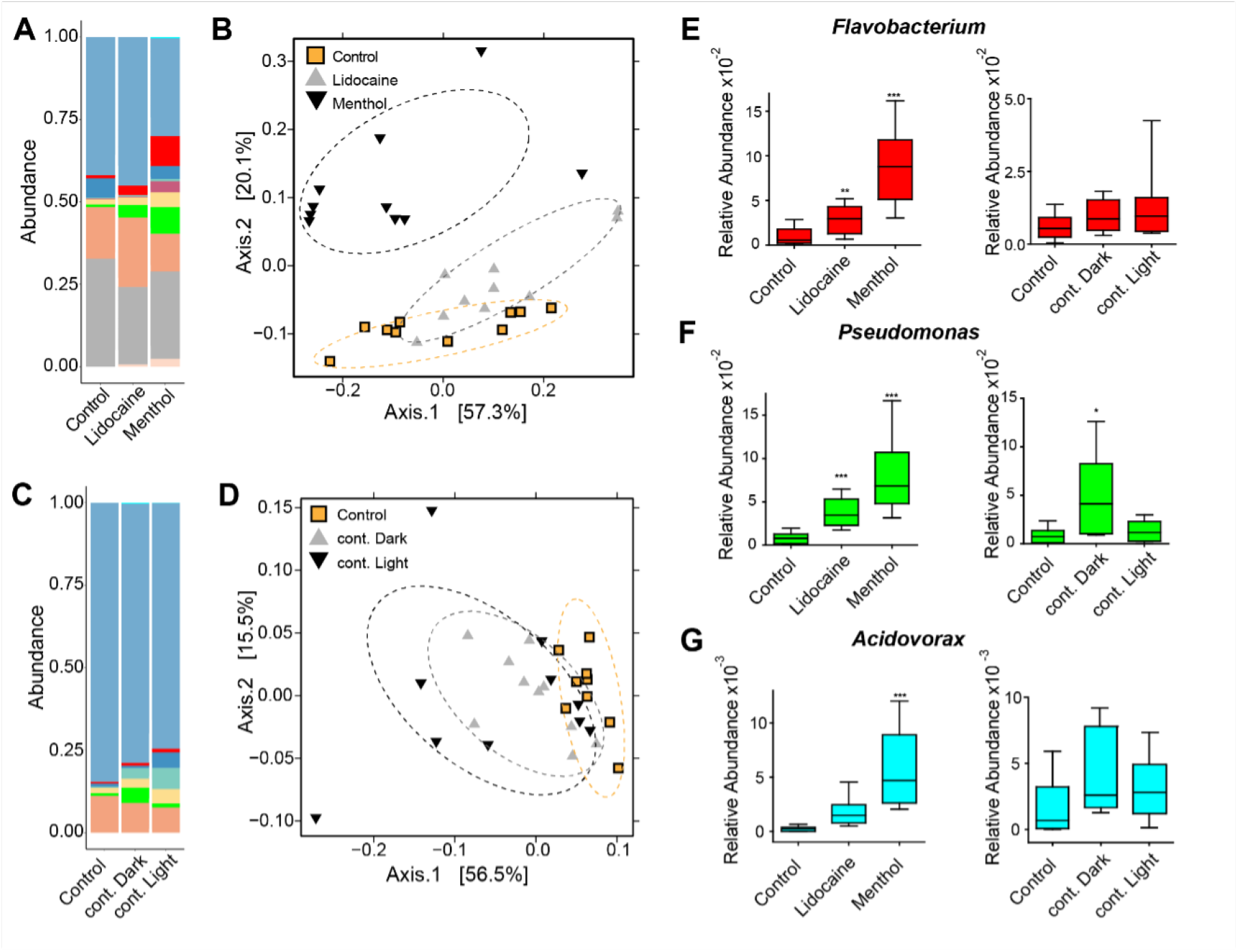
Perturbing the frequency of spontaneous contractions (SCs) over extended time periods shifts the microbial composition in Hydra. Reducing the SC frequency with 48 h treatment of ion channel inhibitors (menthol and lidocaine), continuous light exposure (cont. light), or continuous dark (cont. dark) exposure significantly affects the bacterial community. Control is incubation in freshwater (Hydra medium) and 12 h of light alternating with 12 h of dark conditions. **A**. Bar plot of the relative abundance on the genus level showing the effect of the ion channel inhibitors. **B**. Bar plot of the relative abundance on the genus level showing the effect of the light treatments. **C-D**. Analysis of the bacterial communities associated with the control and disturbed conditions using principal coordinate analysis of the Bray-Curtis distance matrix. The polyps with disturbed contraction frequency show distinct clustering (ellipses added manually). **E-G**. Box plots displaying the fold change of the relative abundance compared to the control of Flavobacterium, Pseudomonas and Acidovorax in response to the different treatments.* *p* ≤ 0.05, * * *p* ≤ 0.01, * * * *p* ≤ 0.001 (ANOVA and Kruskal-Wallis).

### Physics-based model predicts that contractions increase the exchange rate of chemical compounds to and from the surface

We formulated a simple physics-based mathematical model of the transport of chemical compounds to and from Hydra’s surface where the microbes reside. We exploited two key findings from our experimental data. First, between contractions, transport of chemical compounds to and from Hydra’s surface is best described by molecular diffusion rather than fluid advection. Second, PCs are much shorter and faster than IC (*T*_PC_ ≪ *T*_IC_ and *U*_PC_ ≫ *U*_IC_, see Fig. 2C), and each PC causes intermittent shedding of the fluid boundary layer and re-extension of Hydra’s head in a random direction. Thus, each contraction resets the fluid microenvironment and replenishes the chemical concentration around Hydra’s head, while the fluid environment at Hydra’s foot remains always at rest, akin to a permanent IC state.

Our dye visualization and flow quantification showed no noticeable background flows between contraction events (Fig. 2D). The fact that diffusion is dominant between contractions can be formally shown by computing the Péclet number Pe= *LU/D*, which compares the relative importance of advection versus diffusion for the transport of a given compound, such as oxygen. Using the length *L* of Hydra’s body column, the mean flow speed *U* over 1h (averaging over both IC and contracting periods), and the constant oxygen diffusion D at 15° C, we find that Pe is 0.005 ≪ 1, implying that diffusion is dominant between contraction events, even when the flow speed *U* is overestimated by averaging over both IC and SC periods.

To reflect the different fluid microenvironments in the highly motile head and static foot, we approximated the respective head and foot surfaces by two non-interacting spheres of radii *a* and *b* separated by a distance *L* (Fig. 3A, box). When even a few percent of Hydra’s head or foot surface are covered by living microbes or host cells, the diffusion-limited rate of absorption or emission of a chemical compound by these cells is well approximated by a uniformly covered surface [25]. Assuming equivalence of the transport of emitted and absorbed chemical substances (Suppl. Methods), we focus here on absorption only. This implies zero concentration of the chemical compound of interest, say oxygen, at Hydra’s surface. Starting in a compound-rich environment, a concentration boundary layer (CBL) depleted of that chemical compound forms and grows near the surface. In the absence of contractions, the concentration field reaches a steady state with zero rate of change of the compound concentration. Each SC event sheds the chemically depleted fluid boundary layer near the head and effectively resets the depletion zone growth process (Fig. 3A). Accounting for unsteady diffusion following each SC, we computed the growth of the depletion boundary layer over time (stages 1-3 in Fig. 3B), using as an example the diffusion coefficient *D* of oxygen to derive a dimensional timescale *τ* = *a*^2^*/D* (Suppl. Methods). In the absence of fluid boundary layer shedding, such as near Hydra’s static foot, steady state is approached as time increases (stage 4 in Fig. 3B). Near Hydra’s head, however, each SC resets the concentration boundary layer thickness to zero (Fig. 3C, top), resulting in an instantaneous and increased uptake of molecules, such as oxygen, in the head region (blue) compared to the foot (orange). Consecutive contractions increase the maximal cumulative uptake of compounds near Hydra’s head (blue) compared to the foot (orange)(Fig. 3C, bottom).

To investigate the functional implications of experimentally observed temporal distribution of SCs over many hours, we combined our physics-based model of chemical transport (Fig. 3A) with a stochastic model of contraction events. Specifically, Hydra’s SCs follow a Poisson distribution of mean *λ*, for which TIC is described by an exponential distribution of mean 1*/λ* (Suppl. Fig. 2A). We calculated analytically the expected mean and standard deviation of the cumulative uptake *J*_*IC*_ over a single inter-contraction period *T*_IC_ and of the cumulative uptake J over an extended period T (containing multiple *T*_IC_) (Suppl. Methods). We found that the expected mean value of *J*_IC_, given by 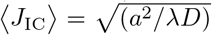, decreases with increasing contraction frequency *λ*, while the expected mean value of J, given by 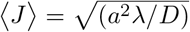, increases with increasing *λ*. This is intuitive: as the contraction frequency increases, the intercontration time *T*_IC_ decreases, so does the expected uptake *J*_IC_ over a single inter-contraction period *T*_IC_. However, the fluid environment gets reset more often, with each resetting event replenishing the chemical concentration, leading to an increase in the expected cumulative uptake *J* over an extended period *T* containing multiple contraction events.

Next, we numerically simulated data sets of inter-contraction intervals *T*_IC_ (Suppl. Fig. 2B) drawn from exponential distributions of mean 1*/λ*, where we let *λ* range from 0 to 10 at 0.05 intervals. For each *λ*, we conducted 10,000 numerical experiments, each lasting for a fixed total time period *T* = 48 h. The obtained number of contraction events in each numerical experiment consisted of one realization taken from a Poisson distribution of mean *λ*. The total number of contraction events over all experiments was normally distributed, as expected from the law of large numbers (Suppl. Methods). We computed numerically the cumulative oxygen uptake J over *T* = 48h for each realization (normalized by the cumulative uptake at steady state), and for each *λ*, we calculated the mean and standard deviation of the computed *J*. Plotting the mean and standard deviation of *J* as a function of *λ* (Fig. 3D), we found that increasing *λ* increases the cumulative uptake *J* and that the mean (solid black line) and standard deviation (thick purple segment) are in excellent agreement with analytical predictions (Suppl. Methods).

SC frequencies on the order of ten CPH and more have been reported in the Hydra [26; 27] indicating that, in theory, increases in uptake rates are possible. In our experience, however, such high frequencies occur only transiently when Hydra has been stressed, e.g., by transferring the animals into very small containers such as concave glass for imaging purposes. At later time points, the contraction frequency of animals in these conditions converge towards an SC rate around 3 CPH (Suppl. Fig. 5), similar to what we observed in our regular setup condition using a large beaker setup (Suppl. Fig. 1A, control condition). This suggests that under unconstrained conditions, Hydra’s baseline CPH is near 3, rather than ten.

We used the model to test the effect of limiting the maximal number of contractions over 48 h to 144 contractions, which is the total number of contractions at an average SC frequency *λ* = 3 CPH. Effectively, this constraint means that once the maximum number of contractions is reached, no additional contractions are permitted during the remainder of the 48 h, mimicking, for example, a finite energy budget for spontaneous contractions. Interestingly, imposing this constraint showed that maximal uptake gains are achieved at contractions frequencies consistent with the imposed constraint (Fig. 3D, thick blue graph), and that increasing *λ* beyond 3 CPH decreases the cumulative uptake.

When testing the effect of a constant-rate contraction activity, i.e., with a constant *T*_IC_ approximately equal to 48*/λ* hours, the model predicts a slightly increased uptake compared to stochastic activity, indicating that a precise rhythm would only confer a small benefit over the stochastic mechanism (Fig. 3D, thin purple and blue graphs). Analogous results hold for the removal for accumulated chemicals produced by microbes at Hydra’s surface (Suppl. Methods).

Our model indicates that Hydra’s spontaneous contractions facilitate a greater exchange rate – including both uptake and release - of chemical compounds in the oral region as compared to the static foot region. When SC frequency is reduced, the maximal exchange rate near the head is reduced as well and, in the extreme case of zero contractions, becomes almost identical to the steady state in the foot region.

The simplicity of the model should not distract from the universality of the mechanism it probes: the effect of stochastic contractions on the transport of chemicals to and from a surface exhibiting spontaneous wall contractions. To make analytical progress, we assumed the surface is spherical, but, by continuity arguments, the conclusions we arrived at are qualitatively valid even for non-spherical surfaces. These conclusions can be restated concisely as follows: fast spontaneous contractions of an otherwise slowly moving surface in a stagnant fluid medium cause impromptu shedding of the fluid boundary layer and lead to improved transport of chemicals to and from the surface between contraction events. Higher contraction frequencies are beneficial, but require additional, may be prohibitive, metabolic cost. A stochastic distribution of contraction events over time, following a Poisson process, produces benefits that are nearly as good as those produced by regular contractions, but without the need for a biological machinery to maintain a precise rhythm. Limiting the total number of contractions leads to decreased performance at contraction frequencies beyond what would allow the contraction events to be Poisson distributed. Taken together, our model suggests that changing the SC frequency will alter the fluid microenvironment and biochemical concentrations experienced by Hydra’s microbial community; in particular, reducing the SC frequency would make the microenvironment near Hydra’s head, where the greatest fluid boundary layer shedding occurs during SCs, more similar to the foot, where minimal fluid boundary layer shedding occurs.

### Reducing the spontaneous contraction frequency changes the colonizing microbiota

To directly probe the impact of SCs on the associated microbial community, we decreased Hydra’s SC frequency by two established methods: 1) continuous exposure to either light or darkness [28; 29] and 2) chemical interference with the ion channel inhibitors menthol and lidocaine [30]. Continuous exposure to light and treatment with ion channel inhibitor lidocaine reduced the SC frequency slightly from an average value of 2.5 CPH in control conditions to 2.3 CPH in continuous light and to 2.0 CPH in lidocaine treatment (Suppl. Fig. 1A,) (note that we were unable to make the measurements in the continuous dark condition). Treatment with ion channel inhibitor menthol almost completely abolished the occurrence of contractions. The treatments reduced SC frequency without significantly changing the frequency of other common fast contractile behaviors, such as somersaulting (Suppl. Fig. 1B and C) [31]. Consistent with earlier studies [28; 30], lidocaine and constant light treatment increased the likelihood of longer TIC; the contraction events remained, however, Poisson-distributed and TIC remained exponentially-distributed (Suppl. Fig. 2A), as assumed in our mathematical model. Computing the average contraction frequency based on the best exponential fit to biological data, we found that it to be equal to 2.5, and 2.2 CPH for the lidocaine, and continuous light treatments, respectively, as opposed to 2.9 CPH for the control, again confirming the reduced contraction rate in the treatment groups.

We next investigated the effects of the altered SC frequency on Hydra’s microbiota. A 48-hour exposure to either pharmacological agents (menthol and lidocaine) or exposure to different light regimes led to a similar and significant shift in the microbial community (Fig. 4, Suppl. Fig. 6). The relative abundance bar plot and Principal Coordinate Analysis (PCoA) of the microbial composition showed a clustering of the different treatments in response to a disturbed contraction frequency (Fig. A-D). The extent of the shift was positively correlated with the magnitude of the decrease in contraction frequency and, consequently, was greatest in menthol treated animals. This shift consisted of a change in the relative abundance rather than a disappearance of bacterial taxa or bacterial load (Suppl. Fig. 8). The fact that both light and pharmacological manipulations resulted in similar effects provide strong evidence that the change in relative bacterial abundance is related to the altered SC frequencies. We further verified by using a minimal inhibitory concentration (MIC) assay that there were no inhibitory effects of the pharmacological substances on the bacteria (Suppl. Fig. 9).

We evaluated the differences between control and treatment groups for up- or down-regulated bacteria using Linear discriminant analysis Effect Size (LEfSe) [32]. When SCs were reduced, the foot specific microbial colonizers Flavobacterium, Pseudomonas and Acidovorax (Fig. 1E-F) became more abundant (Fig. 4E-G). Taken together, these data suggest that a reduced SC frequency alters the microbial composition and potentially expands the biogeography of foot associated bacteria.

Our fluid and transport analysis suggest that these changes in the microbiota result from an altered biochemical microenvironment near Hydra’s head, which is shaped by the SC frequency. In particular, our mathematical model, which faithfully mimics the distribution of ICs under control and disturbed conditions (Suppl. Fig. 2B), predicts that the reduction in SC frequency seen in the experimental treatments (lidocaine, methanol, continuous light) decreases the cumulative uptake or release of any bacteria-relevant chemical compound near Hydra’s head (Fig. 3C), such that the head’s biochemical microenvironment becomes more similar to the static foot region (Fig. 3C).

However, another explanation of the observed changes in the microbiota could be that the contractions are important for the direct displacement of the microbes to other parts of the body. To investigate whether SC have a direct physical impact on the microbiome, we first took video recordings of a normal Hydra polyp colonized with fluorescently labelled bacteria during a full contraction cycle. We could neither observe an overall change in the colonization pattern nor the physical detachment of labelled bacteria (Suppl. Fig 10 and Suppl. Videos 4 and 5). Thus, at short time scale (few seconds), an individual contraction cycle does not affect the biogeography of the microbiome. We also probed whether a lack of recurrent SCs may lead to a remodelled glycocalyx, for example by hindering its redistribution during SCs, and thereby alter the growth conditions of bacteria. Here, we used an antibody specific for a component of Hydra,s glycocalyx [33] and visualized the glycocalyx in animals immobilized for a prolonged (48 h) time period (Suppl. Fig. 11). Since we could not detect major changes in the glycocalyx layer, we concluded that the observed changes in the microbiota are unlikely the cause of glycocalyx remodeling. These results strongly suggest that the fluid boundary layer shedding and resultant chemical exchange by Hydra,s SCs stabilize microbial colonization and contribute to shaping the microbial biogeography along the body column (Fig. 1D-E and Fig. 5A).

## DISCUSSION

The epithelia and microbial symbionts of Hydra’s body walls interface with slow moving or highly viscous fluids. They rely on diffusion for the transport of chemical compounds to and from the surface. They cannot engage in more efficient transport mechanisms via fluid advection [20; 21; 34; 35] because they lack the ciliary or muscular appendages used by other organisms in the low and intermediate Reynolds number regimes to generate unsteady flows and vortices. Our study suggests that Hydra instead facilitates the diffusion process through its spontaneous contractions, which cause transiently and locally higher flow rates and greatly improve the exchange of fluid near the epithelial surface by shedding the so-called fluid boundary layer. This is analogous to a recently discovered “sneezing” mechanism by which marine sponges remove mucus from their surface by recurrent spontaneous contractions [12]. Combining high throughput microbiota profiling, in vivo flow analysis, and mathematical modelling, we found that such exchange facilitates the transport of compounds, including nutrients, waste, antimicrobials, and gases, to and from surface-residing symbiotic microbiota and may take part in maintaining a specific microbiota biogeography along the Hydra’s body column [36]. Interestingly, this distinct spatial colonization pattern along the body column could not only be essential for pathogen defence [3; 13] but could also be important for modulating the frequency of Hydra’s SCs [26], suggesting a feedback loop that could be the focus of future studies.

Our results may also have broader applicability. Hydra’s mucous layer covering the epithelium - an inner layer with stratified organization devoid of bacteria, beneath an outer loose layer colonized by symbionts – is similar to the mammalian colon1. Like Hydra, the peristaltic gut displays recurrent SCs [37; 38; 39] and a viscosity-dominated, low Reynolds number flow regime [40]. Disturbance of this contractile activity (e.g., intestinal dysmotility) is correlated with dysbiosis [41; 42; 43] which can lead to bacterial overgrowth in the small intestine and irritable bowel syndrome [44; 45]. Microfluidic models mimicking the viscous environment and laminar flow of the gut suggest that intestinal contractions mix the luminal content, which appears to modulate microbial density and composition [46]. These observations are consistent with our findings in Hydra and suggest that spontaneous contractions (SCs) might both depend on microbial colonizers and also regulate host-bacteria associations.

Finally, our non-dimensional model enables the generalized prediction that body wall contractions, whether in Hydra or on the gut, cause relative improvement of transport for any compound that is produced or consumed near an epithelial surface in low Reynolds number conditions, thereby simultaneously preventing the build-up of metabolic waste and restoring depleted nutrients. Lowering the frequency of SCs by 10-15%, as seen in our experiments, is predicted to reduce the uptake (or removal) rates of individual compounds by a similar order of magnitude. It is intriguing that such a relatively small change in SC rate altered the microbiome significantly and consistently across treatments. One explanation is that since fluid boundary layer shedding by contractions affects the transport of any chemical compound near the surface, decreasing the contraction rate might alter the uptake or removal of many compounds at once. As shown in a recent study on the response of microbial populations to altered soil properties, such combinatorial effects can greatly exceed, and complicate, the impact of any single factor [47]. Intuitively, and confirmed by our model, increasing the frequency of contractions results in greater transport to and from the surface. Since SCs are energetically expensive processes, Hydra under normal conditions may get along well with a moderate rate of about 3 CPH. Our computational model indicates that this rate still provides significant enhancement over purely diffusive transport, and our experimental data show that lower rates tend to destabilize the microbiome. Taken together, this suggests that Hydra’s rhythm of spontaneous contractions might be operating at a sweet spot of efficiency. Furthermore, we demonstrate that a stochastic distribution of a limited number of contractions over time results in a more effective transport than other strategies, such as alternating periods with a higher frequency of contractions with periods that are static. While contractions at regular time intervals would be slightly more effective still, this would require the presence and maintenance of pacemaker circuits with precisely controlled firing frequency. Though future work is needed, our results suggest that Hydra – and possibly other contractile epithelia systems – achieve near optimal efficiency without the need for a costly high-precision pacemaker system.

Inspired by Dobzhansky,s dictum that “nothing in biology makes sense but in the light of evolution” [48], we speculate that spontaneous contractions were critical early in the evolution of a gut system to maintain a stable microbiota (Fig. 5B). In this context, an observation in a new species of hydrothermal vent Yeti crabs, Kiwa puravida n.sp. is of interest [49]. The crabs farm symbiotic bacteria on the surface of their chelipeds (claws) and wave these chelipeds continuously in a constant rhythm. In search of an explanation the authors stated: “We hypothesize that K. puravida n. sp. waves its chelipeds to shear off boundary layers formed by their epibionts productivity, increasing both the epibionts and, in turn, their own access to food” [49]. Well-controlled perturbation experiments in Hydra may offer a unique model to study both the mechanisms and the in vivo role of spontaneous contractions in more complex systems.

## Acknowledgements

Research in the laboratory of TCGB is supported in part by grants from the Deutsche Forschungsgemeinschaft (DFG), the CRC 1182 “Origin and Function of Metaorganisms” (to TCGB.) and the CRC 1461 “Neurotronics: Bio-Inspired Information Pathways” (Project-ID 434434223 – SFB 1461) (to TCGB and AK). AK is supported by a DFG grant KL3475/2-1. We thank the Central Microscopy Facility at the Biology Department of the University of Kiel for excellent technical support. T.C.G.B. appreciates support from the Canadian Institute for Advanced Research. J.N. and E.K. acknowledge support from the National Institute of Health grant 1 R01 HL 15362201-A1 and the National Science Foundation INSPIRE grant 1608744.

## METHODS and MATERIALS

### Animal manipulation and data analysis

#### Animal culture

Experiments were carried out using *Hydra vulgaris* strain AEP. Animals were maintained under constant environmental conditions, including culture medium (Hydra medium; 0.28 mM CaCl2, 0.33 mM MgSO4, 0.5 mM NaHCO3, and 0.08 mM KCO3), temperature (18°C) and food according to standard procedures [50]. Experimental animals were chosen randomly from clonally growing asexual Hydra cultures. The animals were typically fed three times a week with first instar larvae of Artemia salina; however, they were not fed for 24 h prior to pharmacological interference or light assays, or for 48 h prior to RNA isolation.

#### Pharmacological interference and light assays

To alter the contraction frequency of *H. vulgaris* AEP polyps, the animals were treated for 48 h with either 200 μM menthol (Sigma, Cat. No. 15785) or 100 μM lidocaine (Sigma, Cat. No. L5647), or exposed to 48 h of continuous light or darkness. Control polyps were incubated in Hydra-medium and were exposed to a 12h light-/12h dark cycle over the same time period. Water temperature remained stable at 18°C for all conditions. From hour 24 to hour 32 of the 48h experiment, i.e., for 8h total, ten polyps each were simultaneously video-recorded in a 50 mL glass beaker, using a frame rate of 20 frames per minute. For comparison with prior studies, single polyps were recorded for 8h in a concave slide with a medium volume of 200-500 μL as previously described [26]. Using the video-recordings, we quantified the number of full-body contractions and somersaulting events in ImageJ/Fiji [51] and computed the average contraction frequency per hour.

#### DNA Extraction and 16S rRNA Profiling

To investigate the effect of a reduced contraction frequency on the microbiota, we exposed normal *H. vulgaris* AEP polyps to different pharmacological agents and light conditions over a 48 h time period. Polyps were treated with 200 μM menthol (Sigma, Cat. No. 15785), 100 μM lidocaine (Sigma, Cat. No. L5647), continuous light or continuous darkness for 48 h at 18 °C. Control polyps were incubated in Hydra-medium and were exposed to a 12h light-/12h dark cycle. Afterwards the genomic DNA was extracted from individual polyps with the DNeasy Blood And Tissue Kit (Qiagen) as described in the manufacturer’s protocol. Elution was performed in 50 μl. Extracted DNA was stored at -20°C until sequencing. Prior to sequencing, the variable regions 1 and 2 (V1V2) of the bacterial 16S rRNA genes were amplified according to the previously established protocol using the primers 27F and 338R [52]. For bacterial 16S rRNA profiling, paired-end sequencing of 2 × 300 bp was performed on the Illumina MiSeq platform. The 16S rRNA sequencing raw data are deposited at the SRA and are available under the project ID SRPXXYY. The sequence analysis was conducted using the the DADA2 pipeline in R 3.6.0 [53]. The downstream analysis of the 16S rRNA data (alpha diversity, relative abundance and beta-diversity) was done in R including the packages phyolseq, vegan, DESeq2 and ggplot2 [54; 55; 56; 57]. ASVs with less than 10 reads were removed from the data set. The tables of bacterial abundance were further processed using Linear discriminant analysis effect size (LEfSe) analysis to identify bacterial taxa that account for major differences between microbial communities [32]. An effect size threshold of 3.0 (on a log10 scale) were used for all comparisons discussed in this study. The results of LEfSe analysis were visualized by plotting the phylogenetic distribution of the differentially abundant bacterial taxa on the Ribosomal Database Project (RDP) bacterial taxonomy.

#### Statistics

Statistical analyses were performed using two-tailed Student’s t-test or Mann–Whitney U-test where applicable. If multiple testing was performed, p-values were adjusted using Bonferroni correction.

#### Quantitative real-time PCR analysis (qRT-PCR)

In order to investigate the bacterial community change and validate if there is an increase in the bacterial load, we performed quantitative real-time PCR analysis. The samples from the 16S rRNA profiling experiment were used. Amplification was performed as previously described [58] using GoTaq qPCR Master Mix (Promega, Madison, USA) and specific oligonucleotide primers (EUB 27F, EUB 338R). Four to five biological replicates of each treatment (light, darkness, menthol and lidocaine) and control with two technical replications were analysed. The data were collected using ABI 7300 Real-Time PCR System (Applied Biosystems, Foster City, USA) and analyzed by the conventional ΔΔCt method.

#### Minimal inhibitory concentration assay (MIC)

To test whether the ion channel inhibitors (menthol and lidocaine) have an antimicrobial activity their effect was tested in the MIC assays as previously described [**?**]. The following bacterial strain isolates from the natural *H. vulgaris* strain AEP microbiota were used: *Curvibacter sp*., A*cidovorax sp*., *Pelomonas sp*., *Undibacterium sp*. and *Duganella sp*. [15; 30]. Microdilution susceptibility assays were performed in 96-microtiter well plates. We tested a range of concentrations of the ion channel inhibitors in hydra medium (with an additional 10 % concentration of R2A agar) to match the nutrient poor conditions of the behavioral assay. The concentration range was chosen such that the dose used in the behavioral assay was the middle of the dilution series. The following concentration were tested (in *μ* M): Menthol: 400, 300, 200, 100, and 10 and Lidocaine: 10000, 5000, 2500, 1000 and 100. The inoculum of approximately 100 CFU per well was used. The plates were incubated with the inhibitors for 5-7 days at 18°C. The MIC was determined as the lowest concentration showing the absence of a bacterial cell pellet. The run was designed in a way that every concentration had four replicates.

#### Immunochemical staining of the glycocalyx

To test whether alterations in the contraction frequency has any effect on the glycocalyx of Hydra, we incubated polyps for 48 h in menthol solution or S-medium (control) and visualized the glycocalyx by immunochemical staining and confocal microscopy of whole-mount polyps. To facilitate the detection of the Hydra’s epithelial surface, we used transgenic polyps expressing eGFP in the ectoderm (ecto-GFP line A8 [59]. The glycocalyx was stained using a polyclonal antibody raised in chicken against the PPOD4 protein, generously provided by Prof. Angelika Böttger. The PPOD4 protein has been previously shown to specifically localize to the glycocalyx layer adjacent to the membrane of ectodermal epithelial cells in Hydra [33]. Polyclonal rabbit-anti-GFP antibody (AB3080, Merck) was used to amplify the GFP signal. Immunohistochemical detection was carried out as described previously [58]. Briefly, polyps were relaxed in ice-cold urethane, fixed in 4 % paraformaldehyde, incubated in blocking solution for 1 h, and incubated further with the primary antibodies diluted to 1.0 *μ*g/mL in blocking solution at 4 °C. Following the protocol of Böttger and co-authors, tissue permeabilisation steps were omitted to avoid detection of immature glycocalyx components within epithelial cells. AlexaFluor488-conjugated goat anti-rabbit antibodies (A11034, ThermoFischer) and AlexaFluor546-conjugated goat anti-chicken antibodies (A11040, ThermoFischer) were diluted to 2.0 *μ*g/mL and incubations were carried out for 2 h at room temperature. The samples were mounted into Mowiol supplemented with 1.0 % DABCO antifade (D27802, Sigma). Confocal mid-body optical sections were captured using a Zeiss LSM900 laser scanning confocal microscope. To measure the glycocalyx thickness, the fluorescence profile across the ectoderm has been recorded and quantified for 10 transects, each 10 *μ*m long, using Zen Blue v. 3.4.91 software (Zeiss).

#### GFP labeling of Curvibacter sp. AEP1.3

To visualize the colonization and dynamics during contraction events of Curvibacter sp. AEP1.3, we chromosomally integrated sfGFP behind the glmS-operon via the miniTn7-system as previously established by Wiles et al. 2018 [60]. The protocol was modified in order to manipulate the freshwater bacterium Curvibacter sp. AEP1.3 as followed: Instead of the E. coli SM10 we used the strain E. coli MFDpir as delivery system because bi-and triparental mating was already observed [61; 62]. In addition, the growth medium was changed to the routinely used medium of Curvibacter sp.R2A and antibiotic concentration of the selection media was adjusted to 2 *μ*g/ml.

### Kinematics and fluid flow analysis

#### Fluid boundary layer visualization

The FBL around Hydra was visualized as described previously [63; 19]. Briefly, animals were placed into custom-made cube-shaped acrylic containers filled with Hydra-medium and allowed to acclimatize for 30 minutes. The FBL was visualized using fluorescein dye (Sigma-Aldrich) added near Hydra’s head during intercontraction periods using a transfer pipette. An LED light source was used to illuminate the dye from the side. Videos were recorded using a Sony HDR-SR12 camcorder (1,440 × 1,080 pixels, 30 frames per second; Sony Electronics, San Diego, CA, USA) mounted to a tripod in front of the custom container. Videos were processed to enhance the contrast of the dye with respect to the background using Adobe Premiere Pro (San Jose, CA, USA).

#### Fluid flow analysis

Fluid flow around *H. vulgaris* AEP polyps was quantified using particle image velocimetry under a stereo microscope as described previously [20]. Briefly, *H. vulgaris* AEP polyps were placed in a custom-made acrylic containers with glass walls filled with Hydra-medium that contained 1-μm green-fluorescent microspheres (Invitrogen). A violet laser pointer mounted on a custom micromanipulator stage was aligned with a plano-concave cylindrical lens with a focal length of −4 mm (Thorlabs, Newton, MA, USA) to create an excitation light sheet that captured the particles in a ca. 1 mm thick plane across or along the animal’s body column. The green-emitting particles were recorded from top or side using a stereo microscope equipped with a long pass filter (to remove the violet excitation wavelength) and a camera (same as above) mounted to the eyepiece. Videos were recorded at 60 frames per second. We used Matlab (MathWorks, Natick, MA, USA) with an open-source code package (PIVlab [64]) to measure the particle displacement field across the illuminated plane at each time point and derive the instantaneous planar flow velocity field. From this vector field, the flow velocity magnitude (speed) profiles along lines extending perpendicular form hydra’s surface (Fig. 2E and F) were extracted to determine the FBL thickness (Suppl. Fig. 5), defined here as the distance normal from the surface to the point in the fluid where flow velocity has reached 90% of free stream velocity (here assumed zero), as commonly done in biological systems [17].

#### Body kinematics

From the videos described above, body kinematics were extracted by manually labeling the head and the foot of Hydra in the video and then tracking the displacement of the labels over time using the open-source software Fiji [51] with plugin TrackMate [65]. Using these displacement data, we computed Hydra body length and movement speed using Matlab.

### Physics-based modelling of chemical transport to Hydra’s surface

#### Concentration field surrounding an absorbing surface

We approximated Hydra’s head by a sphere of radius *a* for simplicity, and we analyzed the case in which only molecular diffusion in the suspending fluid governs transport (uptake) of a chemical species of diffusivity *D* to Hydra’s spherical head. Assuming axisymmetry, the concentration of the chemical compound *C*(*r, t*) depends on radial distance *r* from the center of Hydra’s head and *t* is time. The unsteady diffusion equation in spherical coordinates is given by ([66; 25])

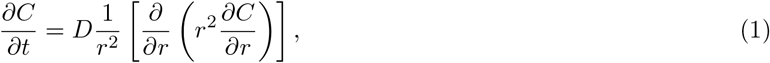

subject to absorbing boundary conditions *C*(*t, r* = *a*) = 0 at Hydra’s surface and constant concentration far from the Hydra *C*(*t, r* → ∞) = *C*_∞_. The unsteady solution is given by

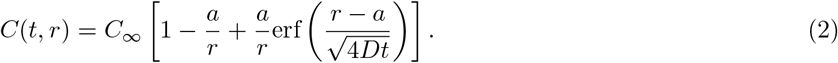

where erf(·) is the error function with erf(0) = 0 and erf(∞) = 1. Note that this solution satisfies the boundary and initial conditions converges as *t* to the steady state solution *C* = *C*_∞_(1 − *a/r*).

The radial concentration gradient is given by

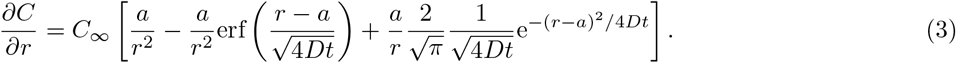

The unsteady uptake of a given chemical compound is defined by *I* = − ∮ **n** *D*▽*C*d*S*, where **n** is the unit normal, and d*S* = 2*πR*^2^ sin *θ*d*θ* is the element of surface area of the sphere,

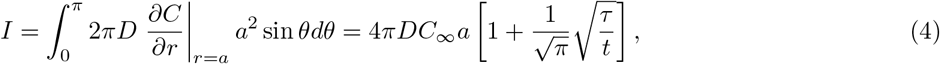

where *τ* = *a*^2^*/D* is the characteristic diffusion time scale.

At steady state, the concentration is given by *C*_*ss*_(*r*) = *C*_∞_(1 − *a/r*); the instantaneous uptake of a chemical compound is *I*_ss_ = 4*πDC*_∞_*a*, and the cumulative uptake *J*_ss_ over a time interval *T* is *J*_ss_ = *I*_ss_*T*.

#### Concentration field surrounding an emitting surface

The transport of a chemical compound of diffusivity *D* produced by emitters covering Hydra’s head is equivalent but opposite to that of a compound absorbed at the surface. To verify this statement, let 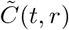 be the concentration field around an emitting surface. The concentration field 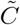 is governed by the unsteady diffusion equation in (1), albeit for a different set of boundary conditions. At the surface, the concentration is given by the boundary condition 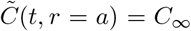, and far from the surface the concentration is nil, 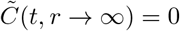. Introducing the transformation 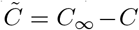, and by linearity of (1), we get that the unsteady solution is 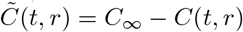, where *C*(*t, r*) is given in (2),

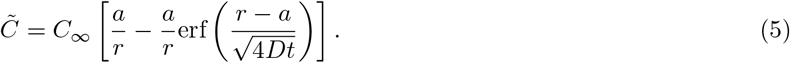

The radial concentration gradient 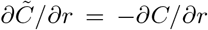 is the opposite of the absorbing gradient in (3), and the unsteady production of a given chemical compound *Ĩ* = − *I* is opposite to the unsteady uptake *I* in (4). It thus suffices to study either chemical absorption or production, with the understanding that the two systems are equivalent. In the following we focus on chemical absorption.

#### Concentration uptake modified by contraction events

Considering a contraction event at *t* = 0 and assuming no additional contractions during an interval *T*_*i*_, corresponding to an intercontraction time period, the cumulative uptake during this time period becomes

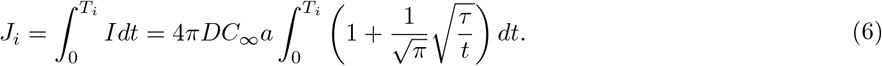

Upon integration, we get

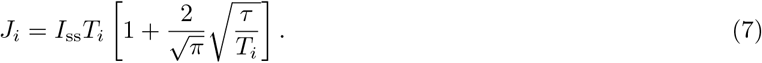

The change in the cumulative intake over one intercontration period *T*_*i*_ relative to the cumulative intake at steady state over the same time period *T*_*i*_ is given by

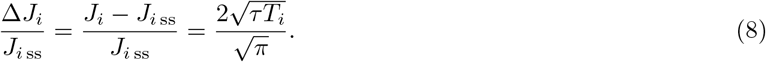

If we had *N* contraction events during a time interval *T*, including the first event at *t* = 0, with inter-contraction time periods *T*_*i*_, *i* = 1, …, *N*, that are not necessarily equal, such that, 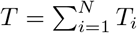, then the total uptake during that period would be (noting that *J*_ss_ = *I*_ss_*T*)

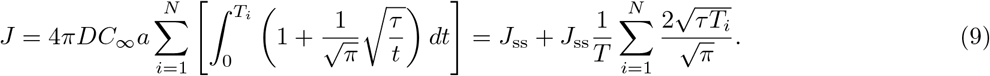

The change in the total cumulative intake with multiple contraction events over the total time period *T* relative to the cumulative intake at steady state over the same time period is given by

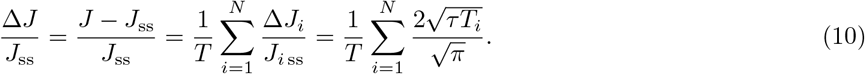

### Coupling chemical transport with stochastic contraction events

We have established in this work that the number of contraction events in a total time period *T* is a Poisson process with expectation *λT* and variance 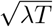, and that the intercontraction time *T*_*i*_ is exponentially distributed with expectation 1*/λ* and variance 1*/λ*. Here, *λ* is the number of contractions per time unit. Here, we derive analytical expressions of the expectations and variances of the incremental and cumulative uptake of a chemical compound between and across contraction events, respectively.

#### Expectation and variance of incremental and cumulative uptake

By definition, the change in cumulative uptake Δ*J*_*i*_*/J*_*i* ss_ over an interval *T*_*i*_, is a function of the intercontraction time *T*_*i*_, which is an exponentially distributed random variable. Therefore Δ*J*_*i*_*/J*_*i* ss_ itself is a random variable. We wish to compute its probability distribution function. Using standard tools from stochastic calculus [67], we arrive at the probability distribution function of the random variable Δ*J*_*i*_*/J*_*i* ss_,

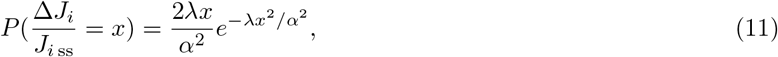

where 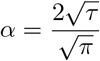 is a constant introduced for notational convenience.

The expectation of the change in incremental uptake over an interval *T*_*i*_ is given by 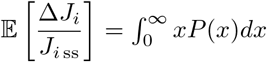. Upon integration by parts, we arrive at

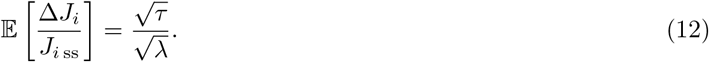

Alternatively, the expectation can be computed directly using the *law of the unconscious statistician* (LOTUS)[67]. The variance is defined as

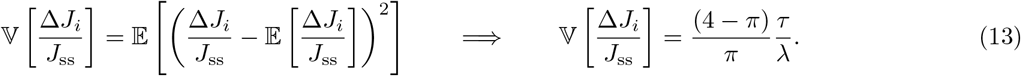

The change Δ*J/J*_ss_ in cumulative uptake over the time interval *T* spanning several intercontraction time periods *T*_*i*_ is also a stochastic random variable defined as the random sum of the random variable Δ*J*_*i*_*/J*_ss_ over a random number *N* of contraction events within *T*, given in (10). While a closed form expression of the probability distribution function of Δ*J/J*_ss_ is not readily available, analytical progress can be made by computing the expectation of Δ*J/J*_ss_

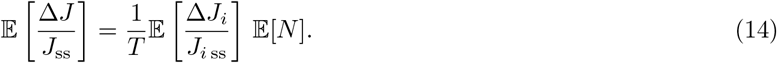

Since contraction events follow a Poisson process, the expectation of *N* contraction events within a time period *T* is given by 𝔼[*N*] = *λT*. Therefore, we can express the expectation of the change in cumulative chemical uptake over an interval of time *T* by

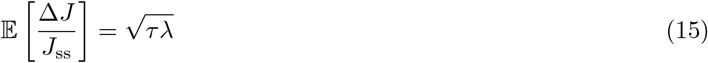

The variance of Δ*J/J*_ss_ conditional on *N* contractions during a fixed interval *T* is expressed by

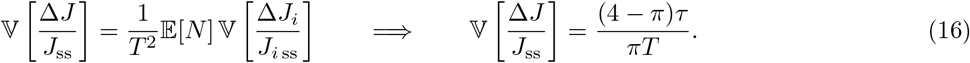

To summarize, all random variables and their corresponding probability distribution function, expectation (mean), and standard deviation (square root of variance) are listed in Table 1.

**Table 1.**
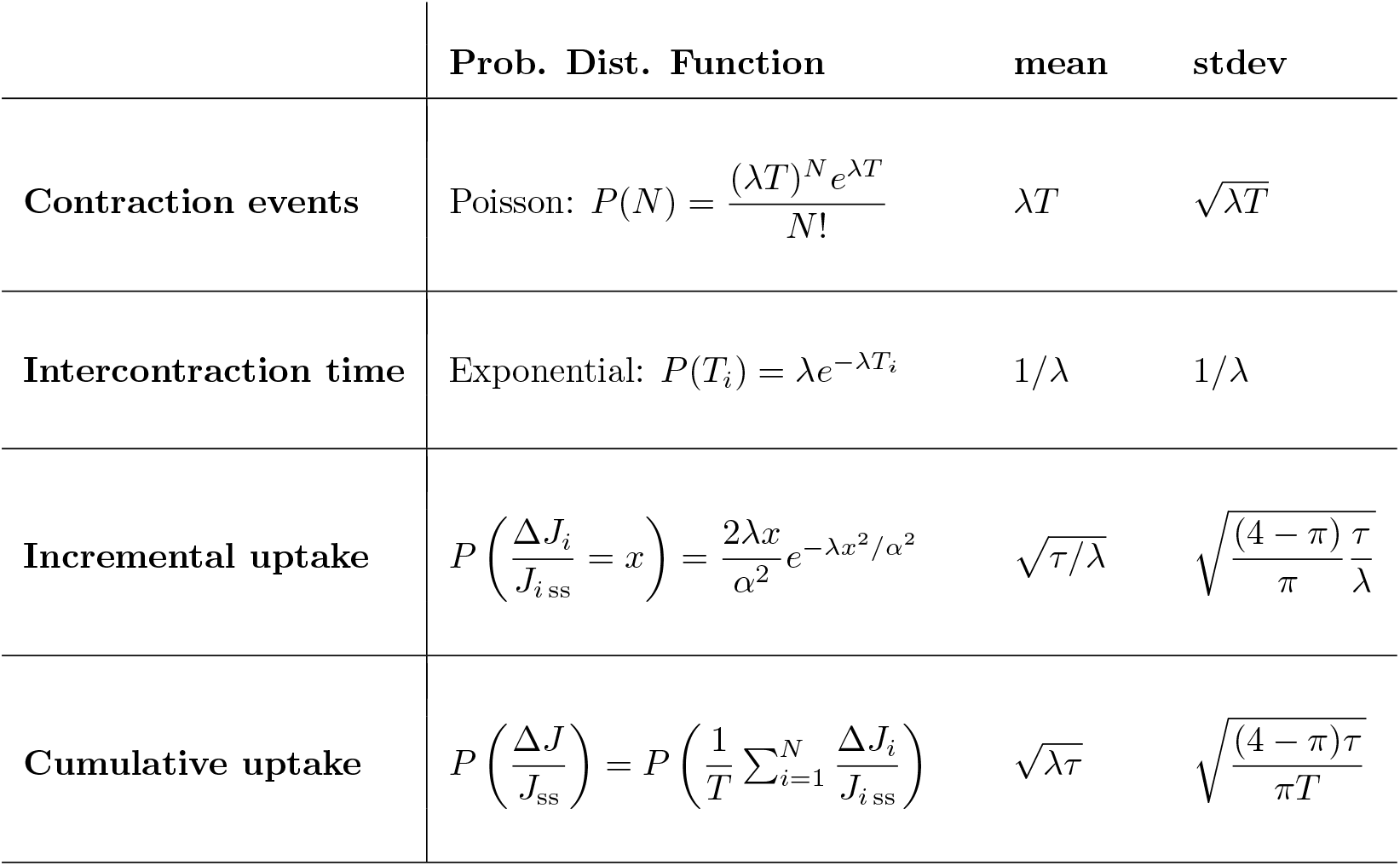
Random variables: The number of contraction events *N* in a time period *T* follows a Poisson distribution, and the intercontraction times *T*_*i*_ follow an exponential distribution. Combining this stochastic description of contraction events and intercontraction times with physics-based models of chemical uptake over the interval *T*, we arrive at a stochastic model of incremental uptake over each intercontraction period *T*_*i*_ and cumulative uptake over the interval 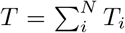. The p.d.f., mean (expectation) and standard deviation (stdev = square root of the variance) of the input (*N, T*_*i*_) and output 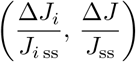 random variables are listed here.

#### Numerical simulations

In addition to these analytical calculations of the expectation and variance of the incremental and cumulative uptake, i.e., Δ*J*_*i*_*/J*_*i* ss_ over *T*_*i*_ and Δ*J/J*_ss_ over *T* = *T*_*i*_, respectively, we also conducted numerical simulations of contraction events and intercontraction times. We used a common algorithm that exploits the fact that intercontraction times are exponentially distributed, and we simulated intercontraction times until the maximum time period *T* is achieved. We checked the histograms, expectation, and variance of both contraction events and intercontraction times obtained from the numerical simulations and verified that they are all in line with theory, with nearly machine precision error in the expectation values obtained from simulations and theory for the range of *λ* we examined.

For each value of *λ*, we conducted 10,000 experiments over a time interval *T* = 48 h, and for each experiment, we computed Δ*J*_*i*_*/J*_*i* ss_ and Δ*J/J*_ss_. We then calculated the expectation (mean) and variance (square of the standard deviation) of Δ*J/J*_ss_ over all 10,000 experiments. Results are shown in main manuscript as a function of *λ* and are in-agreement with the theoretical predications in Table 1.

