## Supplemental Figures and Video Captions for "Spontaneous body wall contractions stabilize the fluid microenvironment that shapes host-microbe associations"

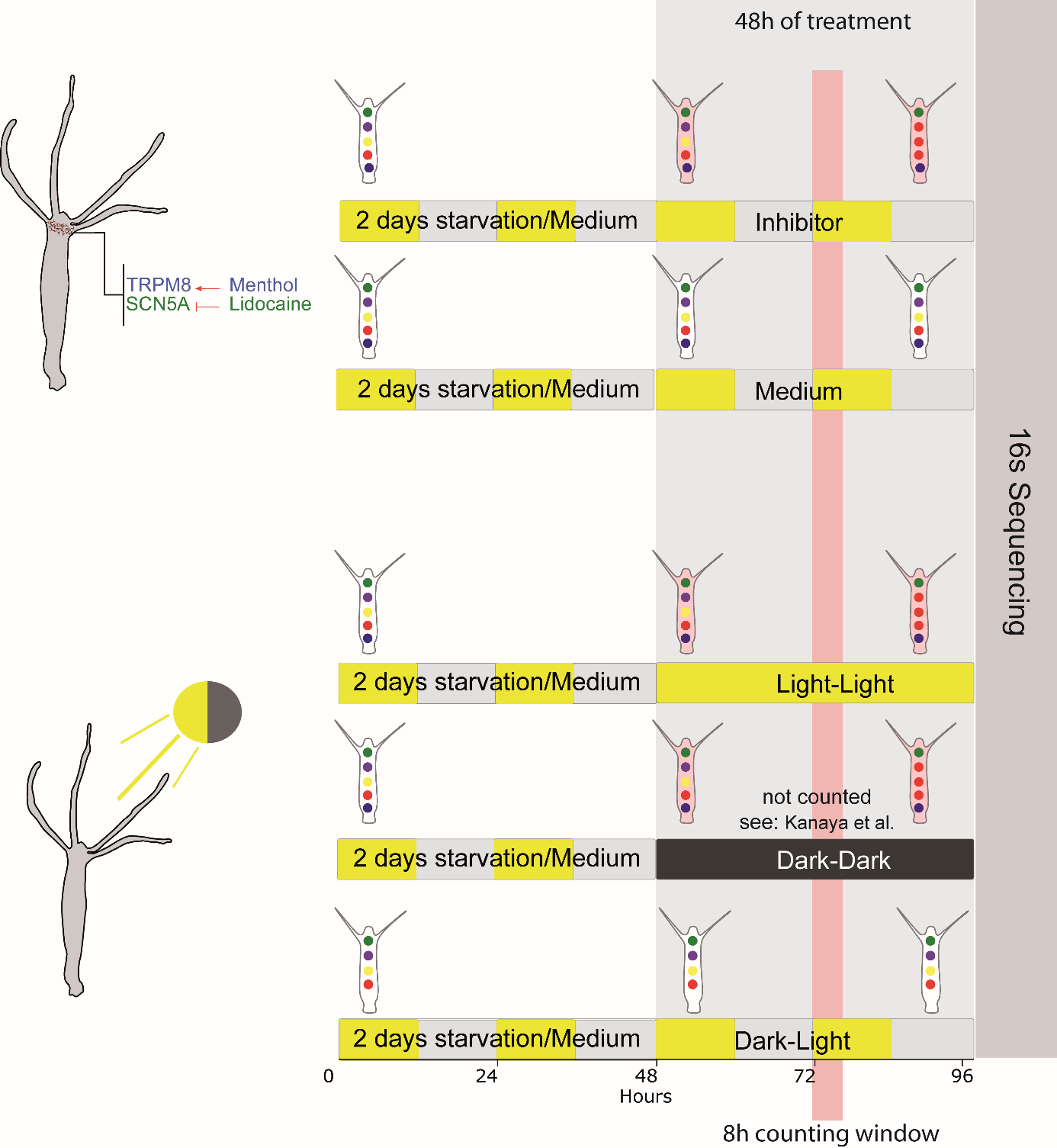

**B**

**A**

**Supplementary Fig. 1. The experimental setup for identifying the effect of spontaneous contraction on the microbiota**. We used two different approaches to identify the effect of reduced spontaneous contractions on the microbiota. **A.** Use of ion channel inhibitors, which reduce spontaneous contractions. Animals were incubated for 48h at 18°C/80% humidity with medium containing either lidocaine or menthol, or with medium alone (control). Highlighted in red is the 8h time window when contractions were counted. After the 48h of treatment, the gDNA was extracted from the polyps and send for 16S rDNA sequencing. **B**. The contractile behavior of hydra is also light sensitive and was reduced under light-light and dark-dark conditions. Contraction data and gDNA were collected as above.

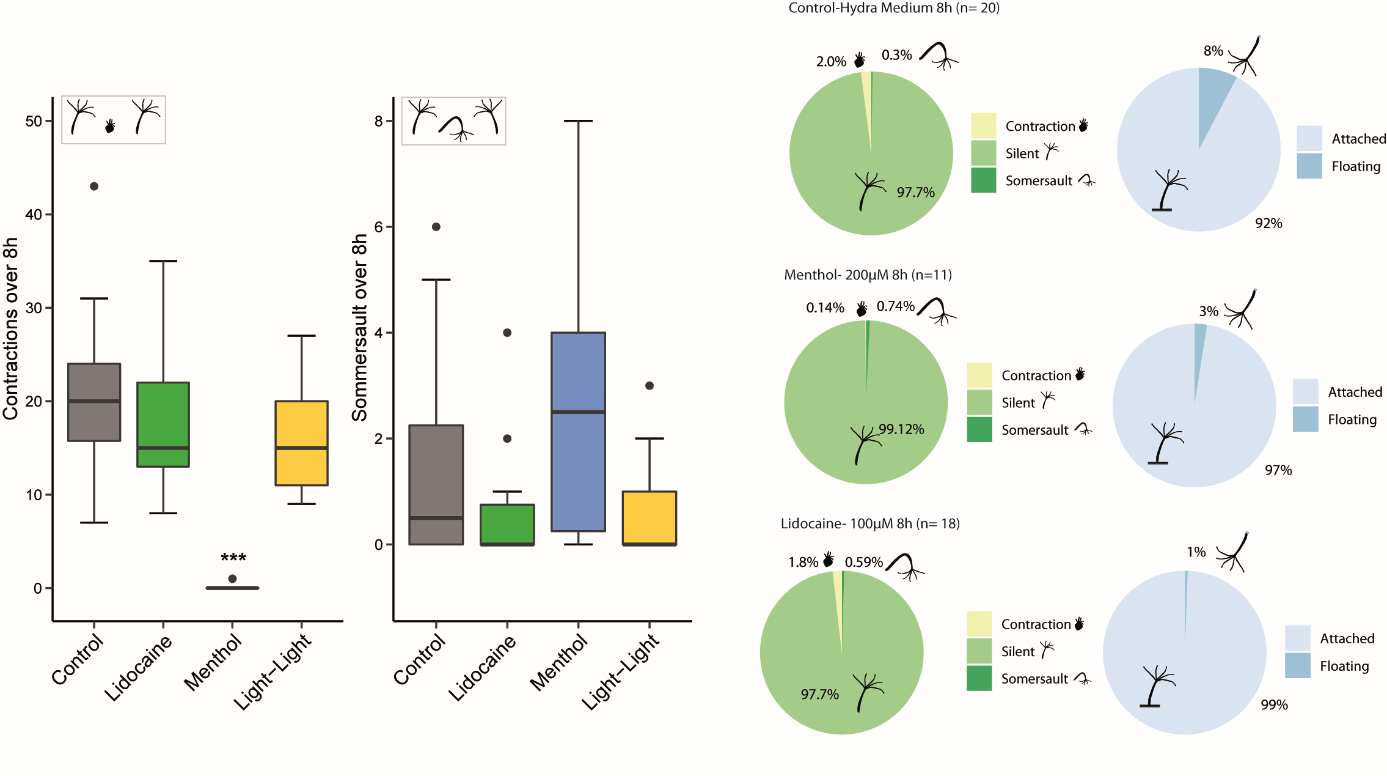
**Supplementary Fig. 2. Behavioral analysis of hydra over 8h**. **A and B**. Quantification of the number of contractile behavior events over time. **A**. The number of spontaneous contractions is reduced under the experimental treatment conditions compared to control. Menthol effectively abolishes all spontaneous contractions whereas the other treatments are reducing the frequency. **B**. Hydra somersault behavior is not affected by the different treatments and remains a rare event. **C.** Percentage of time different Hydra behaviors occurred. The time a polyp spent in a spontaneous contraction event was reduced by the lidocaine and menthol treatment. In addition, time of floating at the surface is reduced in the menthol and lidocaine treatments compared to the control.

**C**

**B**

**A**

**Supplementary Fig. 3.** **Spontaneous contractions are modeled mathematically as a Poisson process.**  **A.** The temporal distribution of spontaneous contraction events is shown for all treatment groups in a density plot (top) and a spike raster plot (bottom). **B** Characteristic for a Poisson-like process, the time measured between contractions (T_IC_) follows an exponential distribution for all conditions (except menthol), with Lidocaine and Light-Light treatments characterized by a longer tail compared to control, reflecting lower average contraction frequencies. Shown is the best exponential distribution fit. **C** Combined histogram density plot of the experimental T_IC_ data in all conditions (left) compared to a typical simulated data set (right) that recapitulates key experimental observations, particularly the longer tails of the treatment groups (here we show just one data set per condition, and each data set contains 10,000 randomly sampled intervals).

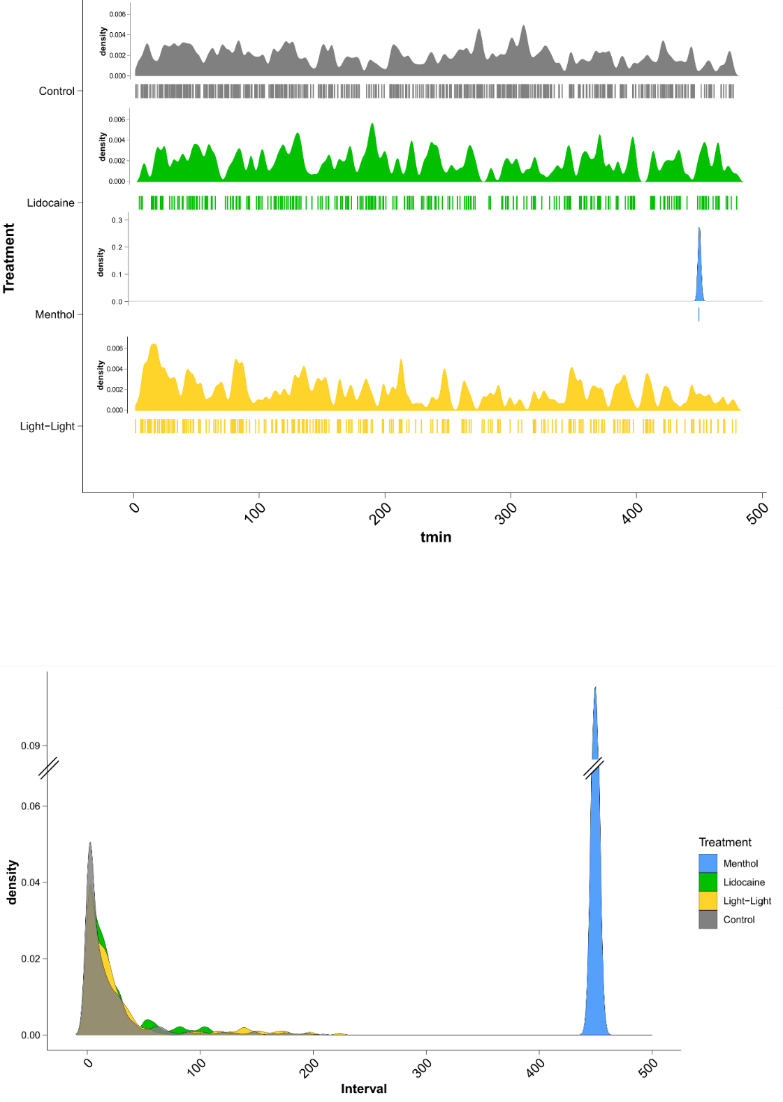

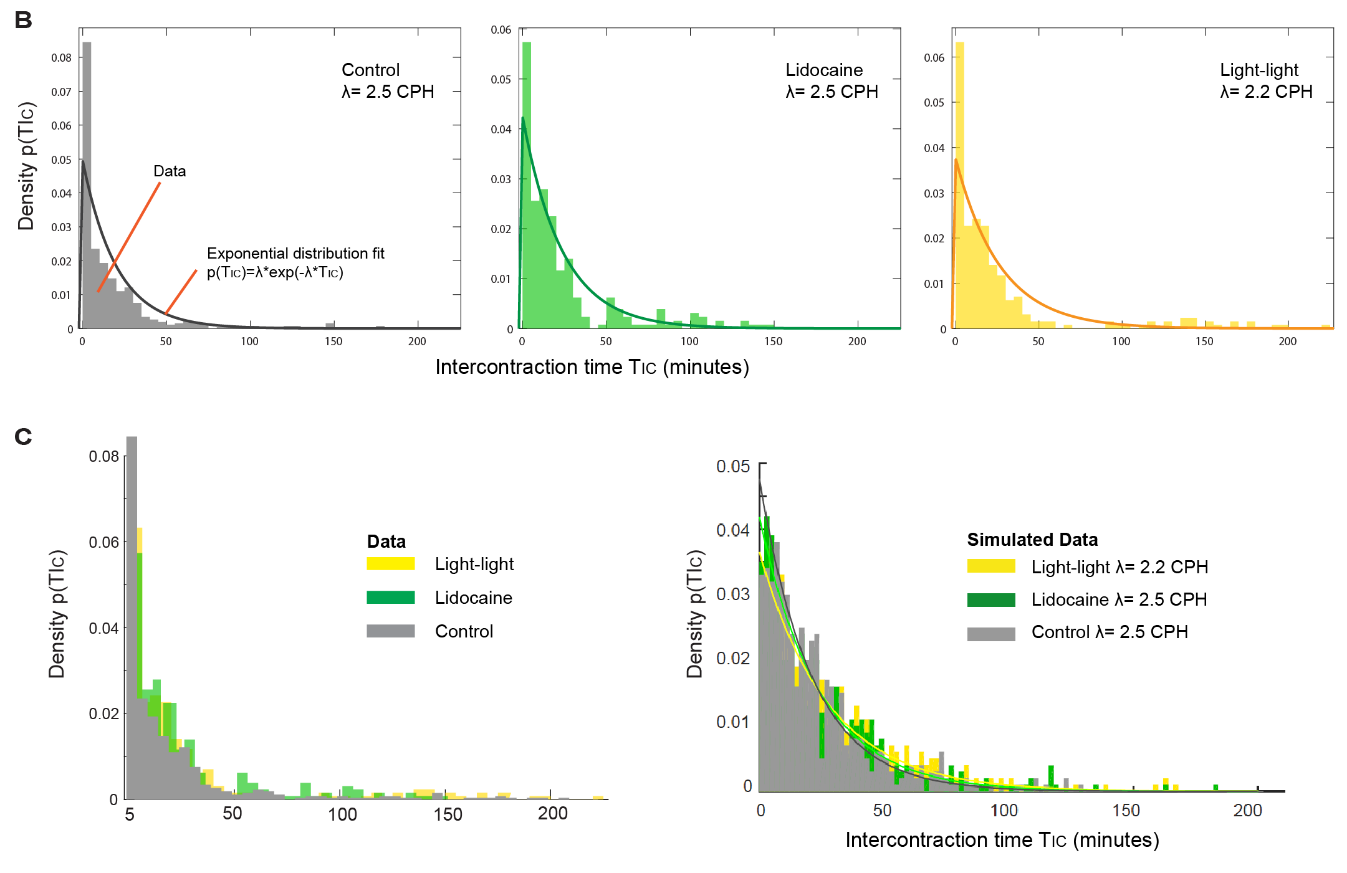

**Time (minutes)**

**A**

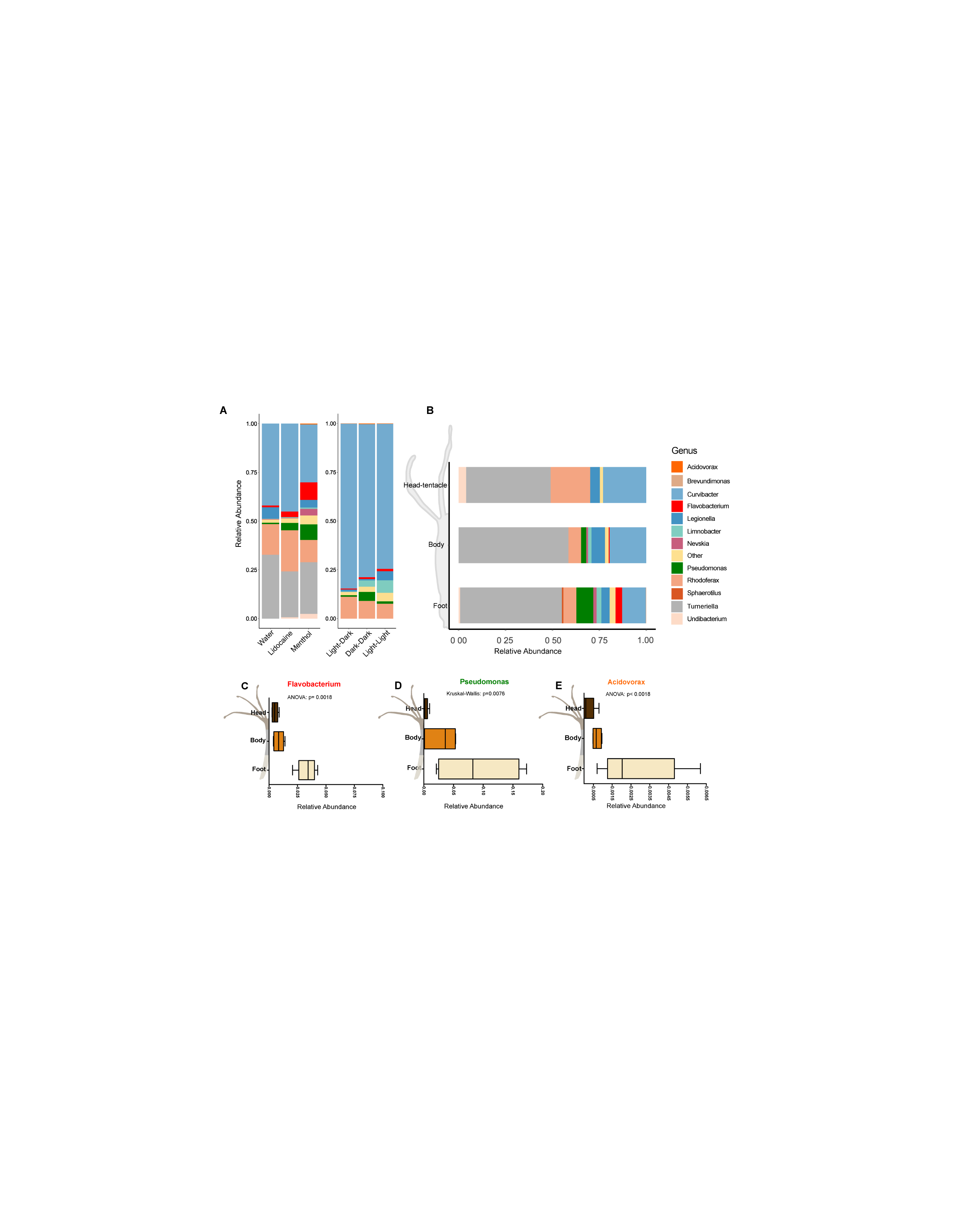

**Supplementary Fig. 4. The microbiota changes due to reduced contraction frequency**. **A**. The relative abundance of the microbiota after 48h of treatment shows that lidocaine/menthol and dark-dark/light-light treatments cause shifts in microbiota composition. *Flavobacterium*, *Pseudomonas* and *Acidovorax* increased in their relative abundance in the treatment groups. **B**. The spatial distribution of the microbiota along the body axis under normal/undisturbed condition. **C-E**. The spatial distribution of the genera that are increasing in abundance after treatment. All three have a higher abundance in the foot region.

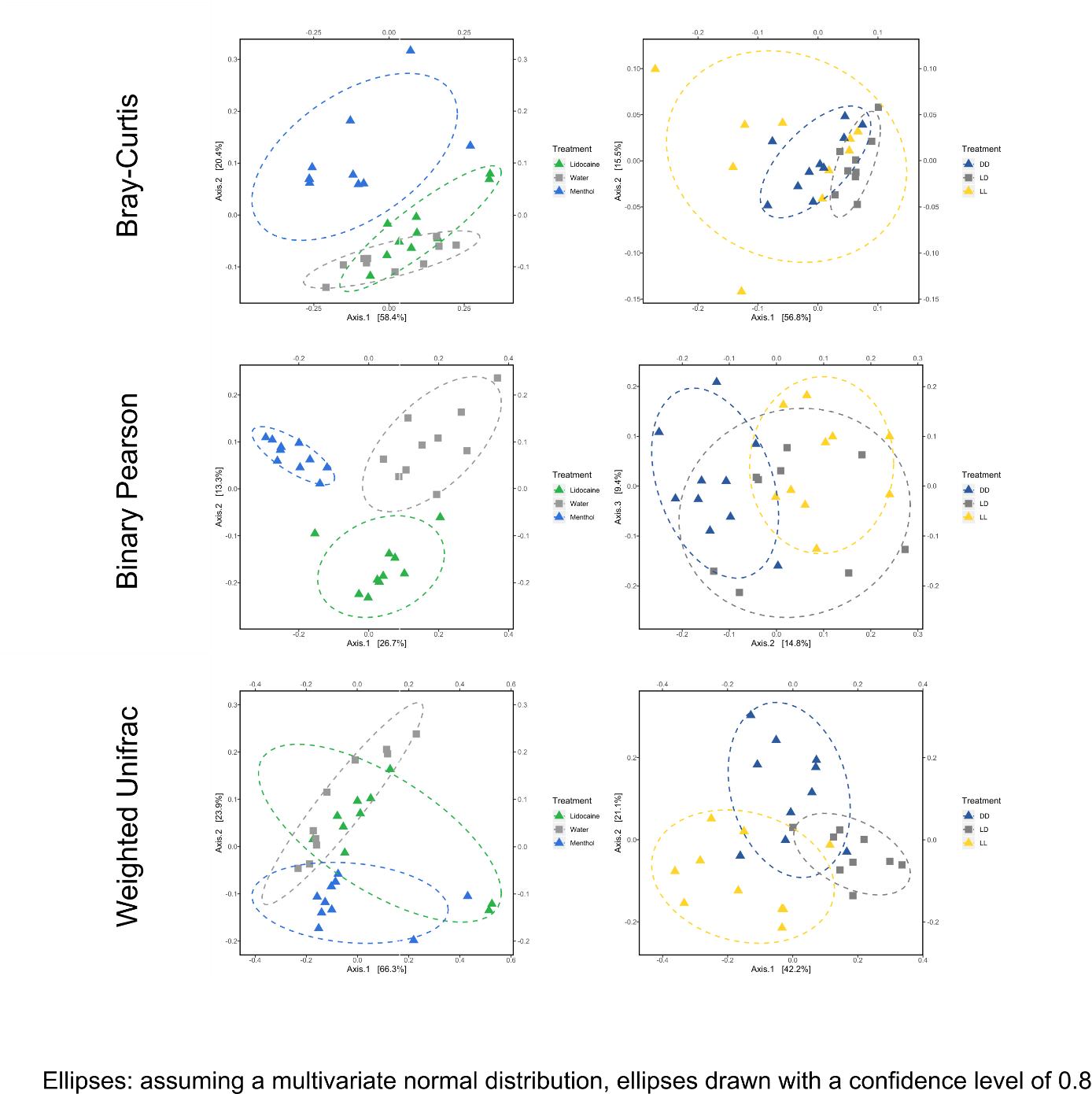

**A**

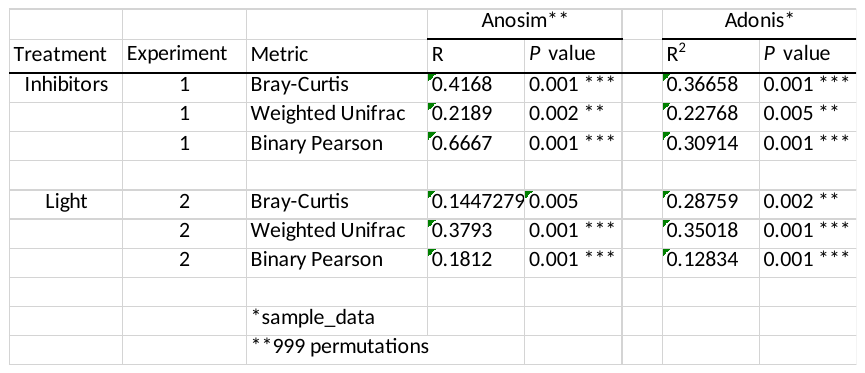

**Supplementary Fig. 5. Analysis of the bacterial communities using different distance matrixes. A.** Different principal coordinate analyses (PCoA) of the bacterial communities using different distance matrixes. Here: Bray-Curtis, Binary Pearson and Weighted Unifrac. In all three PCoA plots, a shift of the bacterial communities is visible in respond to the different treatments. **Table**. Statistical analysis of the PCoAs using Anosim and Adonis. Experiment-1 and -2 are two independent experiments conducted in an identical way using either the ion channel inhibitor menthol and lidocaine or different light conditions.

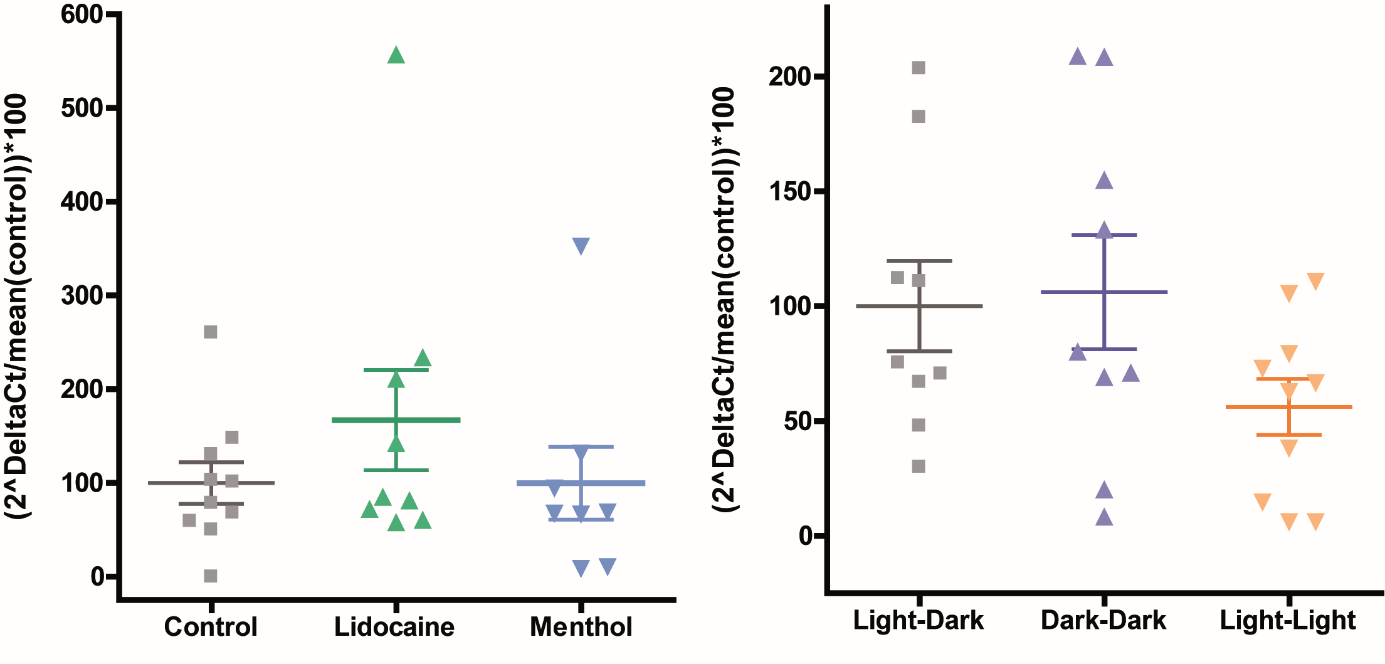

**Supplementary Fig. 6.** **The bacterial load is not affected by ion channel inhibitors or light treatments.** The bacterial load does not show a significant difference compared to control (Kruskal-Wallis (non-parametric) and Dunn’s multiple comparison test) tested by qRT-PCR using Eub-primer (bacterial load measure) and Elon-F (gDNA host measure). The data consist of two technical replicates (two runs) and 4-5 biological replicates. The error bars show the mean with SEM. The data are normalized to the mean of the respective control and shown as percentages.

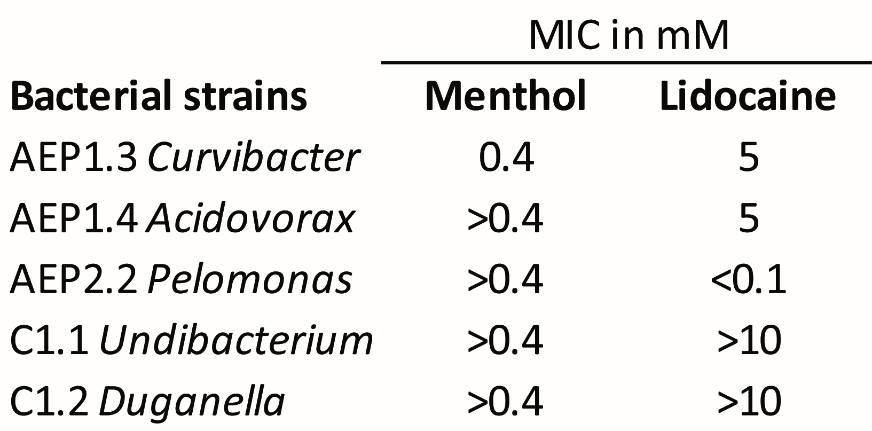

**Supplementary Fig. 7. The minimal inhibitory concentration of the ion channel inhibitor**. Neither ion channel inhibitors showed any antimicrobial effect in the concentration range used in the behavioral assays (Menthol 200 µM, Lidocaine 100 µM). The effect was evaluated after three days of incubation in the same conditions as the behavioral experiments (18°C/80% humidity/12h-light cycle). (n= 3 per concentration)

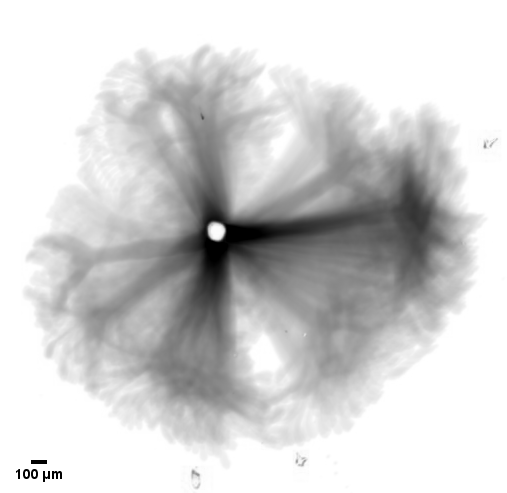

**Supplementary Fig. 8.** The 2D space covered by Hydra’s resting motion and re-extensions in the period of 9 contractions is revealed by this overlay of detected motion. The light spot in the center demarks the foot, which stayed fixed.

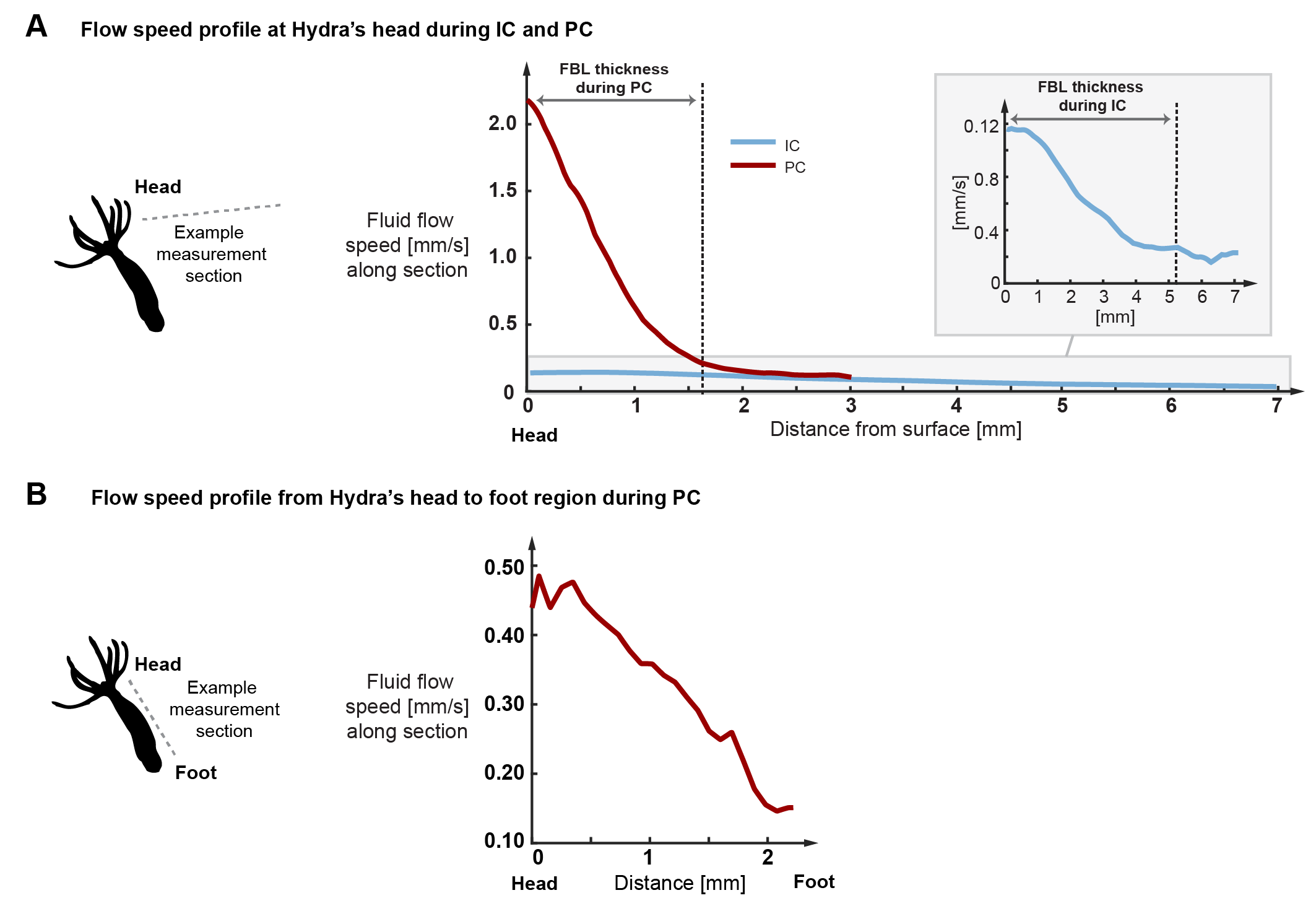

**Supplementary Fig. 9.** Typical flow speed profiles derived from the PIV data. **A.** Flow speed profiles during IC and PC measured along the sections shown in Fig. 2E and F starting at hydra’s head and ending in the free stream flow. As indicated, FBL thickness is defined as the distance at which the flow speed decreases to approximately 10% of the surface speed. **B.** Flow speed profile during PC measured along the body column from head towards the foot region. Towards the foot, the flow declines rapidly to a fraction of maximal speed.

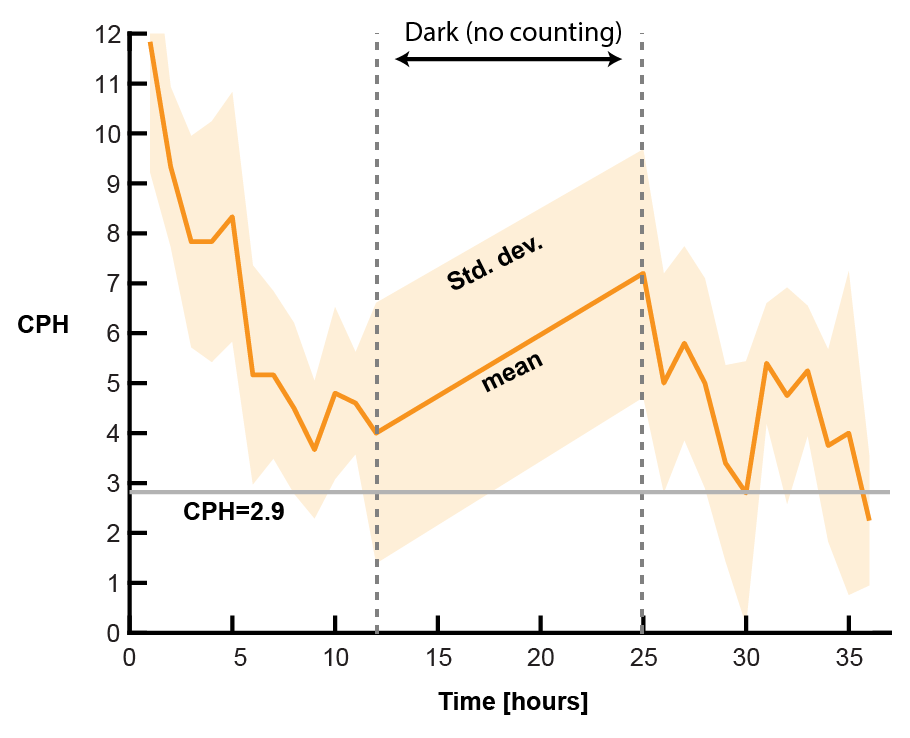

**Supplementary Fig. 10.** Spontaneous contraction frequencies of Hydra when placed in small liquid volumes can reach up to 12 CPH. Over time, these rates decrease substantially and converge to 3 CPH or less, comparable to the baseline rate observed in our control conditions where animals were placed in a large fluid volume.

**Suppl. Video 1:** *Hydra* alternating between rest (intercontraction time) and spontaneous contractions.

**Suppl. Video 2:** Dye visualization of fluid boundary layer (FBL) dynamics during rest (intercontraction time), when it remains stable, and during a spontaneous contraction, when much of the FBL is shed.

**Suppl. Video 3:** Particle image velocimetry to quantify fluid boundary layer dynamics during rest (intercontraction time) and spontaneous contractions.

**Suppl. Video 4:** *Hydras* tentacles during a full body contraction colonized with fluorescent *Curvibacter sp.* AEP1.3 (white signal on the surface).

**Suppl. Video 5:** *Hydras* body column during a full body contraction colonized with fluorescent *Curvibacter sp.* AEP1.3 (white signal on the surface).
